# 3D spatial transcriptomics reveals the molecular structure of input and output pathways in the mouse olfactory bulb

**DOI:** 10.1101/2025.02.19.639192

**Authors:** Nell Klimpert, Mihaly Kollo, David H. Brann, Chichun Tan, David Barry, Ying Ma, Andreas T. Schaefer, Alexander Fleischmann

## Abstract

A core organizing principle of the vertebrate brain is its symmetry along multiple axes. However, the precision with which neurons, circuit modules, and brain regions align to these axes remains poorly understood. Here, we used 3D spatial transcriptomics to reconstruct the anatomical and molecular organization of the mouse olfactory bulb. We mapped the positions of nearly one thousand molecularly distinct glomeruli, the structural and functional units of odor processing, revealing highly symmetric organization across hemispheres. Within each bulb, we delineated a curved axis of symmetry that divides pairs of sister glomeruli. Gene expression in the olfactory epithelium predicted glomerular position with near-glomerular resolution. However, glomerular symmetry did not extend to deeper layer mitral and granule cells, suggesting a reorganization from sensory input to cortical output pathways. Our findings provide the first comprehensive map of the olfactory bulb and reveal how its molecular structure is instructed by epithelial gene expression programs.

## INTRODUCTION

Brains are highly variable in size, structure, and complexity across the animal kingdom, yet a fundamental shared feature across all vertebrate brains is bilateral symmetry: brain regions are duplicated along the midline, resulting in mirror-symmetric left and right hemispheres. Furthermore, even within hemispheres, brain areas often exhibit mirror-symmetric maps, for example in auditory and visual cortex^1,2^. Despite such overall symmetry, many structural and functional asymmetries have been described. Cortical areas involved in auditory processing and vocalization in humans and non-human primates are larger in the left hemisphere^3–5^. In birds including pigeons and chickens, visual pathways are dominant in the left hemisphere^6,7^, and in the zebrafish brain, visual and olfactory information processing is lateralized in structurally asymmetric habenulae^8,9^. At what resolution, then, are brains symmetric? Limitations in the spatial resolution of fMRI make it difficult to identify differences in the human brain beyond regional asymmetries. In the smaller brains of invertebrates, resolving hemispheric symmetry at the cellular level is possible; however, cellular resolution for mammalian brains with millions of neurons is currently beyond reach. The olfactory system provides a unique opportunity to probe symmetry at the resolution of independent structural and functional units: glomeruli in the mouse olfactory bulb (OB) represent segregated channels of information processing that are approximately 100 µm in diameter.

Olfactory sensory neurons (OSN) in the olfactory epithelium (OE) project axons to glomeruli in the OB. Odor-evoked activity in OSNs is relayed to OB projection neurons, mitral and tufted cells, which form microcircuits with deep-layer inhibitory granule cells to route information to the olfactory cortex (**Fig. 1A**). OSNs expressing one of approximately 1,100 odorant receptor (OR) genes^10,11^ converge their axons to ∼2 sister glomeruli per OB hemisphere^12–18^. Prior work has described mirror-symmetry across OB hemispheres, which may be functionally relevant for stereo-sampling odor information via inter-hemispheric connections through the anterior olfactory nucleus^13,19,20^. In addition, mirror-symmetry exists within each OB hemisphere^21,22^, with each OSN type believed to innervate one medial and one lateral sister glomerulus. However, whether this overall symmetry reflects a broad compartmentalization of the OB or the precise, mirrored arrangement of individual glomeruli remains unknown. Moreover, it is not known whether these duplicated glomerular inputs align with molecularly distinct output domains.

**Figure 1:**
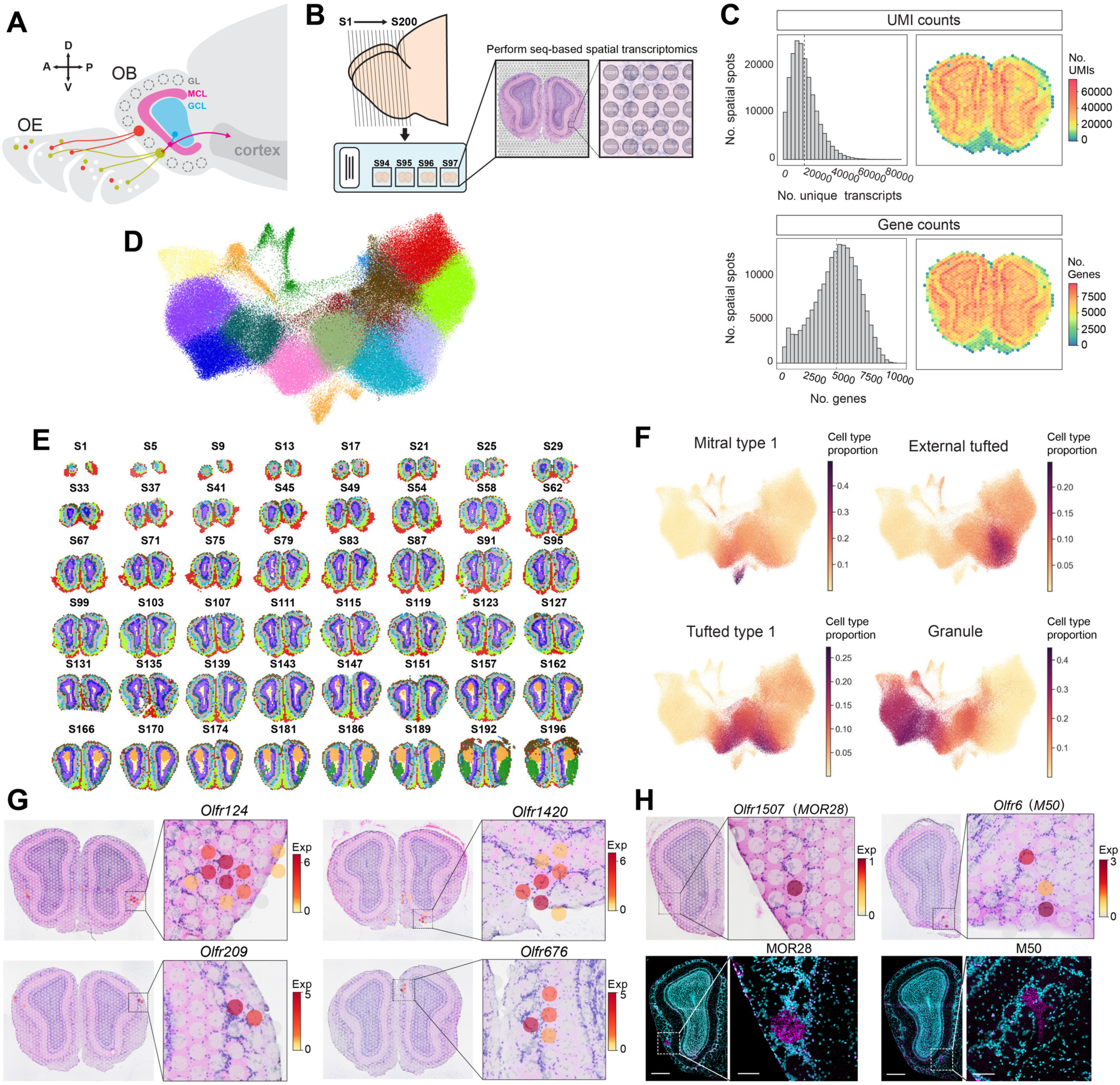
Generation of a comprehensive molecular map of mouse olfactory bulb using sequencing-based spatial transcriptomics. **(A)** Schematic representation of olfactory circuit. Molecularly distinct olfactory sensory neurons in the olfactory epithelium project axons to glomeruli in the olfactory bulb. Olfactory information is then routed to the olfactory cortex via mitral cell projection neurons and shaped by interneurons such as inhibitory granule cells. OE, olfactory epithelium; OB, olfactory bulb; GL, glomerular layer; MCL, mitral cell layer; GCL, granule cell layer. **(B)** Schematic of spatial transcriptomics experiment. 200 consecutive sections of OB tissue were mounted onto capture probe microarrays. Sections were stained with hematoxylin and eosin and imaged, then tissue was permeabilized to allow mRNA capture. 200 individual libraries were prepared, providing spatially localized gene expression data. **(C)** RNA quality control. **Top:** Histogram of the average numbers of unique transcripts, or unique molecular identifiers (UMIs) per capture spots. On average, 13,941 UMIs were captured per capture spot across all 200 tissue sections; heatmap shows a representative image of transcript numbers in a single section. **Bottom:** Average numbers of genes per capture spot. On average, 4,547 genes were captured per capture spot across 200 sections; heatmap shows a representative image of the number of genes in a single section. **(D)** Leiden clustering of 190,445 integrated capture spots across 200 tissue sections. **(E)** Capture spots colored by Leiden cluster in 48 representative tissue sections in the dataset, revealing clustering based on OB layers. **(F)** Identification of capture spot cell type identities by deconvolution of spatial transcriptomics data with reference OB scRNA-seq. Color represents cell type proportion in capture spots. **(G)** Examples of gene expression of *Olfr124*, *Olfr676*, *Olfr1420*, and *Olfr209* in single tissue sections. OR expression is restricted to 3-8 capture spots surrounding a single glomerulus. **(H)** Validation of OR expression. Left: Capture of *Olfr1507* transcripts via spatial transcriptomics in section 135 (top) aligns with targeting of MOR28 glomerulus (magenta) in adjacent section (bottom), localized via immunohistochemistry (see Methods). Right: Capture of *Olfr6* transcripts in section 142 (top) aligns with targeting of M50 glomerulus (magenta) in adjacent section (bottom). DAPI in cyan. For bottom: Left scale bar = 500µm; right scale bar = 100µm.

Here, we generated a comprehensive, three-dimensional molecular map of the mouse OB. We performed sequencing-based spatial transcriptomics on 200 consecutive tissue sections of OB and captured OR transcripts within glomeruli to confidently define the positions of 968 glomeruli. We characterize the mirror-symmetric organization of glomerular maps across hemispheres and identify lateral and medial domains separating sister glomeruli within each OB. Using single-cell gene expression data from OSNs, we uncover genetic signatures that can predict glomerular position with high accuracy. Finally, we show that the symmetry of the glomerular input layer is restructured in the mitral and granule cell output layers.

## RESULTS

### 3D spatial transcriptomics of the mouse olfactory bulb

To generate a comprehensive molecular map of the mouse OB, we used sequencing-based spatial transcriptomics (Visium, 10x Genomics) to capture mRNA transcripts from serial sections of the OB (200 cryosections, 10 µm thick) (see Methods, **Fig. 1B**, **Fig. S1**). The technology utilizes printed microarrays with 55 µm diameter mRNA capture spots to integrate gene expression and spatial information from individual tissue sections. We initially assessed the quality of our RNA sequencing data by integrating data across all sections. Our dataset contained 190,445 capture spots with, on average, 4,547 unique genes and 13,941 unique transcripts per capture spot (**Fig. 1C**). Next, we clustered the capture spots based on their transcriptomic similarity using the Leiden algorithm and deconvolved these clusters with an established snRNA-seq reference dataset of mouse OB to evaluate their cellular composition (see Methods, **Fig. 1D and E**, **Fig. S2A and B**). We found that capture spots clustered in a layer- and cell-type-specific manner (**Fig. 1E and F**). For example, spots in the mitral cell layer captured high levels of the known mitral and tufted cell marker *Cdhr1*^23–25^, spots in the glomerular layer contained high levels of the external tufted cell marker *Ly6g6e*^25^, and spots in the outer olfactory nerve layer contained the known OSN marker *Omp*^26,27^ (**Fig. S2C**).

To comprehensively determine glomerular identity, we leveraged the fact that OR mRNA transcripts are present in OSN axon termini^14,16–18^. We found that, in single tissue sections, individual OR genes were typically expressed in localized regions in the glomerular layer. For example, *Olfr124*, *Olfr1420*, *Olfr209*, and *Olfr676* were each expressed in 3-8 capture spots centered over individual glomeruli, with minimal background expression (**Fig. 1G**, **Fig. S3**). OR transcript capture was validated using immunohistochemistry on a small set of saved tissue sections, where expression profiles of *Olfr6* and *Olfr1507* were localized within M50 and MOR28 glomeruli, respectively, in neighboring sections (**Fig 1H**). Furthermore, expression of a single OR gene was consistently detected within the same glomerulus in consecutive sections. For example, expression of *Olfr3* was restricted to 4-7 capture spots surrounding a single visually identifiable medial glomerulus across 8 consecutive sections (**Fig. 2A**).

**Figure 2:**
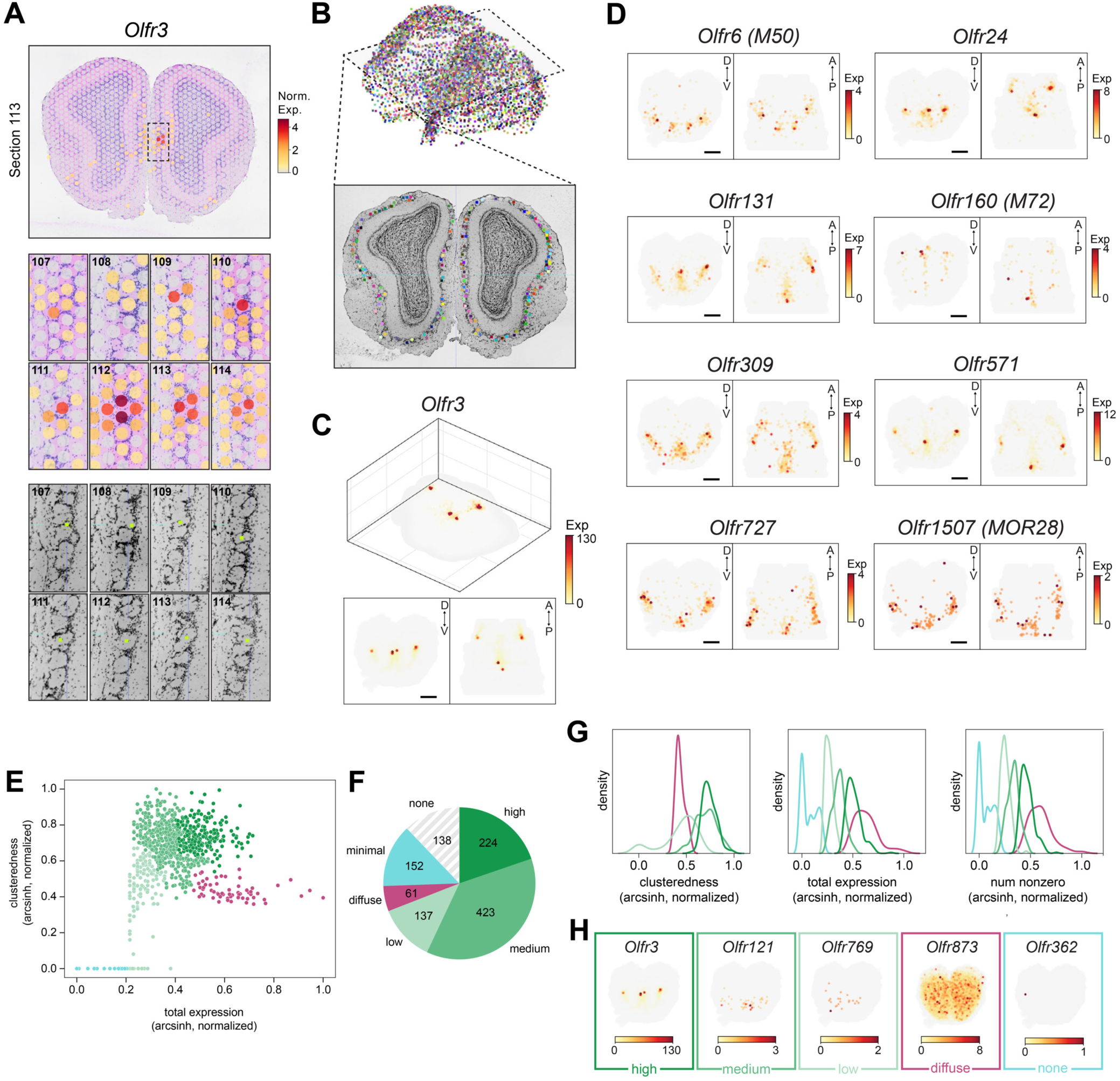
Morphological alignment and transformation of capture spots in 3D. **(A)** Top: Normalized expression of *Olfr3* is overlayed against H&E image. High normalized expression is observed in capture spots surrounding a single glomerulus across consecutive sections. **Bottom:** The center of a single glomerulus is annotated across 8 consecutive sections. Glomerular annotations were used to refine the alignment of the glomerular layer. **(B)** 3D visualization of the 2550 manually annotated, histologically defined glomeruli, with each colored cube representing the center of a single glomerulus. Bottom shows the coronal view of a single section. **(C)** Raw expression of *Olfr3* visualized in 3D (top) and as a maximum Z-projection along the anterior-to-posterior axis and Y-projection along the dorsal-to-ventral axis (bottom). 3D representation of OR expression reveals defined areas of high expression consistent with glomeruli. **(D)** Raw expression of 9 OR genes, visualized as a maximum Z-projection (left) and Y-projection (right). Examples show areas of high expression (red) vs. low expression (yellow) and illustrate variability in expression across OR genes. **(E)** Classification of gene expression patterns based on clusteredness of capture spots expressing the OR and total expression across the tissue. Clusteredness is determined as the silhouette score of clusters after a 3D k-means clustering (4 clusters) of spots. **(F)** Expression patterns across the 1135 OR genes. Expression was detected in 82% of genes with variable expression levels and clusteredness, 5% showed diffuse, non-clustered expression, and 12% were not detected. **(G)** Distribution of descriptors (clusteredness, total expression, number of nonzero capture spots) across different expression categories. **(H)** Example raw data showing the Z-projection of a single example OR gene from each cluster.

To integrate gene expression and spatial information in 3D, we combined manual annotation of over 2,500 glomeruli based on the distribution of nuclei in the reference histology images with computer vision to align tissue sections (**Fig. 2B, Fig. S4**) (see Methods). We then transformed the capture spot coordinates and corresponding gene expression data to align with this morphological reconstruction. We found that sites of high OR expression aligned with morphologically identified glomeruli in 3D (**Fig. 2C and D**). For example, expression of *Olfr3* was localized to 4 restricted sites across both bulbs, consistent with one lateral and one medial glomerulus per bulb (**Fig. 2C**). Moreover, low levels of OR expression often extended from focal regions of high expression in glomeruli to the outer nerve layer, consistent with localization to OSN axons (**Fig. 2C and D**). Overall, our dataset contains OR gene expression data from 997 out of 1,135 annotated OR genes. OR genes varied widely in abundance, likely due to variable expression and capture efficiency^28,29^. We categorized these expression patterns based on the strength of OR expression and parameters describing spatial distribution (**Fig. 2E-H**). Out of the 997 OR genes captured in the dataset, 784 showed expression with a structured and spatially clustered distribution of spot counts with variable expression levels (“Low”, “Medium”, and “High”), and 152 showed minimal expression. The remaining 61 OR genes showed low levels of background expression throughout the tissue volume (“Diffuse”). Taken together, we have generated the first 3D spatial transcriptomics data set with comprehensive gene expression information, including OR expression localized to individual glomeruli, across the entire mouse OB.

### A computational model to quantify glomerular identity and position

We next used a Bayesian framework to quantify glomerular position. We modeled the distribution of OR gene expression in capture spots within a given glomerulus based on glomerular geometry, assuming that the detected expression is linearly dependent on the total overlap between the glomerular volume and the detected spots (see Methods, **Fig. 3B**). We generated predicted count values in each capture spot for any given set of putative glomeruli defined by their position, radius, and expression level. We then optimized the model by iteratively updating the distributions of these glomerular parameters, comparing them to observed OR gene expression using a Markov Chain Monte Carlo (MCMC) algorithm (see Methods, **Fig. 3C and S5**). The resulting Probabilistic Glomerulus Position (PGP) model generated putative locations with different levels of probabilities for 906 ORs. We then confirmed glomerular identity for glomeruli that were assigned a high probability by the model using human expert inspection of predicted positions and raw gene expression data (see Methods, **Fig. S6A and B**). After expert curation, the PGP model yielded a glomerular map containing the quantified positions of glomeruli expressing 233 OR genes, yielding a map of 673 glomeruli (**Fig. 3A**).

**Figure 3:**
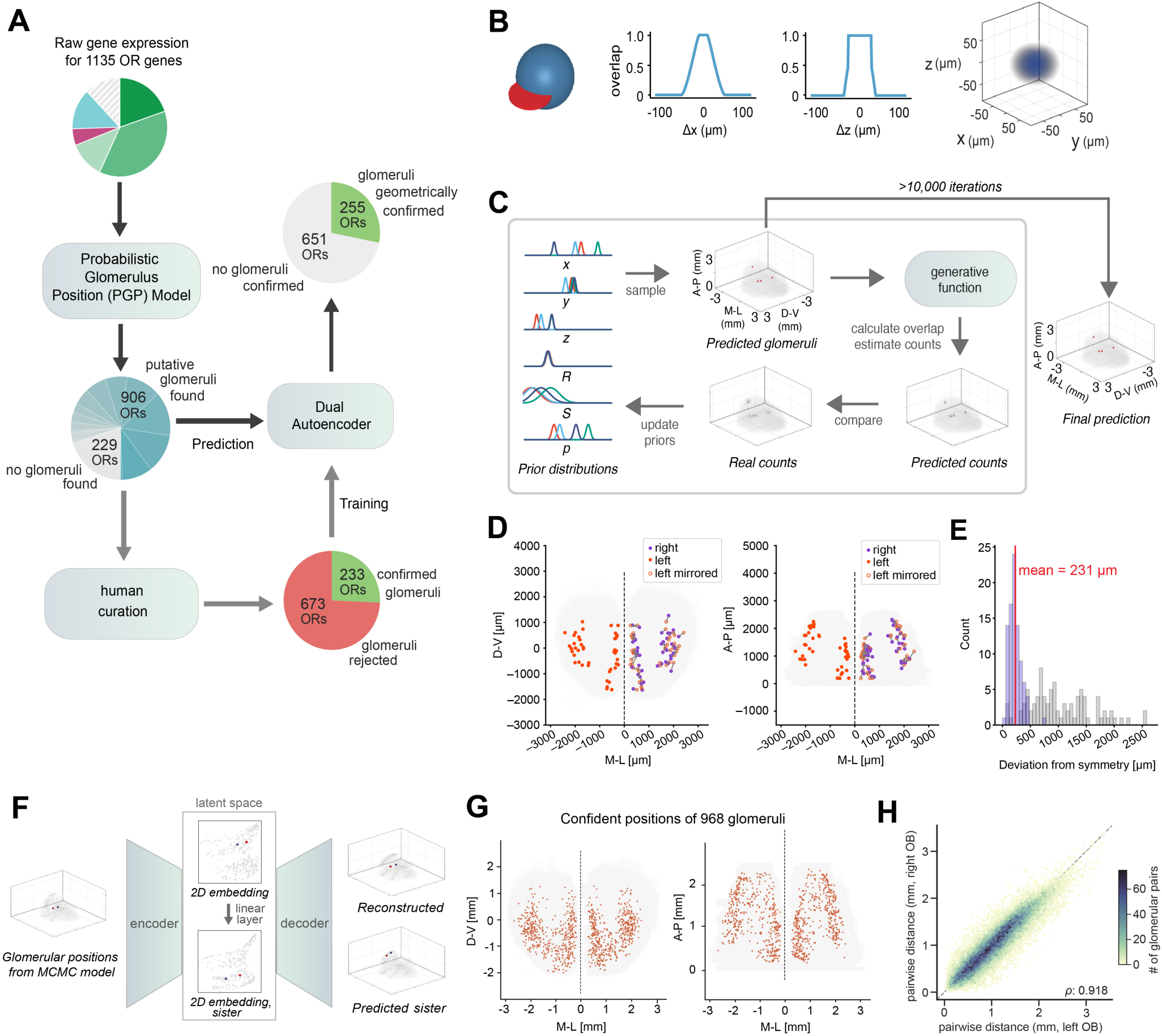
Spatially aware probabilistic modelling to quantify positions of 968 glomeruli. **(A)** Schematic of the process used to position glomeruli from OR gene expression. Initial identification of putative glomeruli was done using a probabilistic spatial gene expression PGP model of count values in proximity to glomeruli. OR genes were categorized based on differences in expression patterns (see also Fig. 2E**-H**). The PGP model fit putative glomeruli at various confidence levels (the hue of the blue segments represents p=<0.1 to =1 for the glomeruli of 906 ORs). A curated dataset of confirmed glomeruli was then defined by human experts (see Methods), which was used to train a dual autoencoder (AE) model to predict the location of sister glomeruli of each putative glomerulus in the unsupervised output of the PGP model. **(B)** The generative function used by the PGP model. Count values at each spot depend on the overlap between the spherical glomerulus and the disk-shaped volume above the spot, resulting in a “squished ball” shaped distribution of count values around the glomerulus position (right). **(C)** The PGP model performs probabilistic inference of glomerulus position (x, y, z), radius (R), strength (S), and probability of validity (p). Samples are drawn from the prior distributions to generate putative glomeruli. The generative function is used to estimate counts, which are compared with real counts to update priors iteratively. A final prediction of glomerular position is determined after greater than 10,000 iterations. **(D)** Left and right bulbs were aligned based on the positions of 233 glomeruli identified by the PGP model and confirmed by expert curation. The resulting left and right glomerular maps show high symmetry, with a close spatial alignment corresponding left-side and right-side glomeruli. **(E)** The resulting left and right glomerular maps show high symmetry, with a mean deviation of 231 µm between corresponding glomeruli. Gray bars represent the hemispheric distance relative to the closest glomerulus of a random OR gene. **(F)** An autoencoder (AE) model predicts the location of sister glomeruli for each putative glomerulus identified by the PGP model. 3D glomerular positions are projected onto a 2D embedding space. A single linear layer is used to predict the 2D latent embedding of sister glomeruli from the original glomeruli. The same decoder is used to decode the positions into final 3D sister glomerulus positions. **(G)** Glomerular map showing the positions of 968 glomeruli expressing 255 OR genes. Glomeruli identified by the PGP model and obeying relative sister geometry defined by the AE model were included in the final map. Left panel shows the Z projection of the glomerular map, and right panel shows the Y projection. **(H)** Linear correlation of pairwise distances between glomeruli in the left OB (x-axis) and right OB (y-axis). Color indicates the number of glomerular pairs.

Visual inspection of these glomeruli revealed two axes of symmetry: glomeruli expressing the same OR were bilaterally mirror symmetric across hemispheres and within each hemisphere. We first compared the glomerular maps across hemispheres to quantify symmetry. We aligned the geometric positions of the two hemispheres to counteract relative rotation and distortions during tissue processing. We then measured the distance between corresponding glomeruli of the same molecular identity from the two OB (**Fig. 3D and E**). Our analysis revealed a remarkably high upper bound of inter-bulbar symmetry, with a mean deviation of 231 µm from perfect symmetry (**Fig. 3E**).

Next, we assessed the symmetry of the glomerular map within each OB hemisphere. Considering the complex, curved three-dimensional geometry of the OB glomerular layer, we used a dual autoencoder (AE) model to describe the relative spatial arrangement of sister glomeruli expressing the same OR. This algorithm uses a two-layer encoder network to map the positions of glomeruli based on the PGP model into a two-dimensional embedding, then reconstructs glomerular positions across bulbs and sister glomerulus positions within each bulb using two identical decoder networks (see Methods, **Fig. 3F**, **Fig. S7 A-F**). The resulting encoding revealed that sister glomeruli were well separated by vector direction in the latent space (**Fig. 4A and B**), corresponding to medial (red) and lateral (blue) glomeruli. This allowed us to define an intra-hemispheric axis of symmetry and assign a medial and lateral label to each glomerulus based on its relative position to its sisters in an unsupervised manner (**Fig. 4C-F**). Finally, we utilized the predicted positions of sister glomeruli proposed by the autoencoder trained on curated glomeruli to confirm the validity of the entire set of putative glomeruli generated by the PGP model (see Methods, **Fig. 3A**). The resulting glomerular map is comprised of 968 high-confidence glomeruli expressing 255 OR genes (**Fig. 3G**, **Fig. S8**). The final glomerular map exhibited a high degree of symmetry across OB, with pairwise distances of glomerular positions well correlated with each other between left and right OB (**Fig. 3H**).

**Figure 4:**
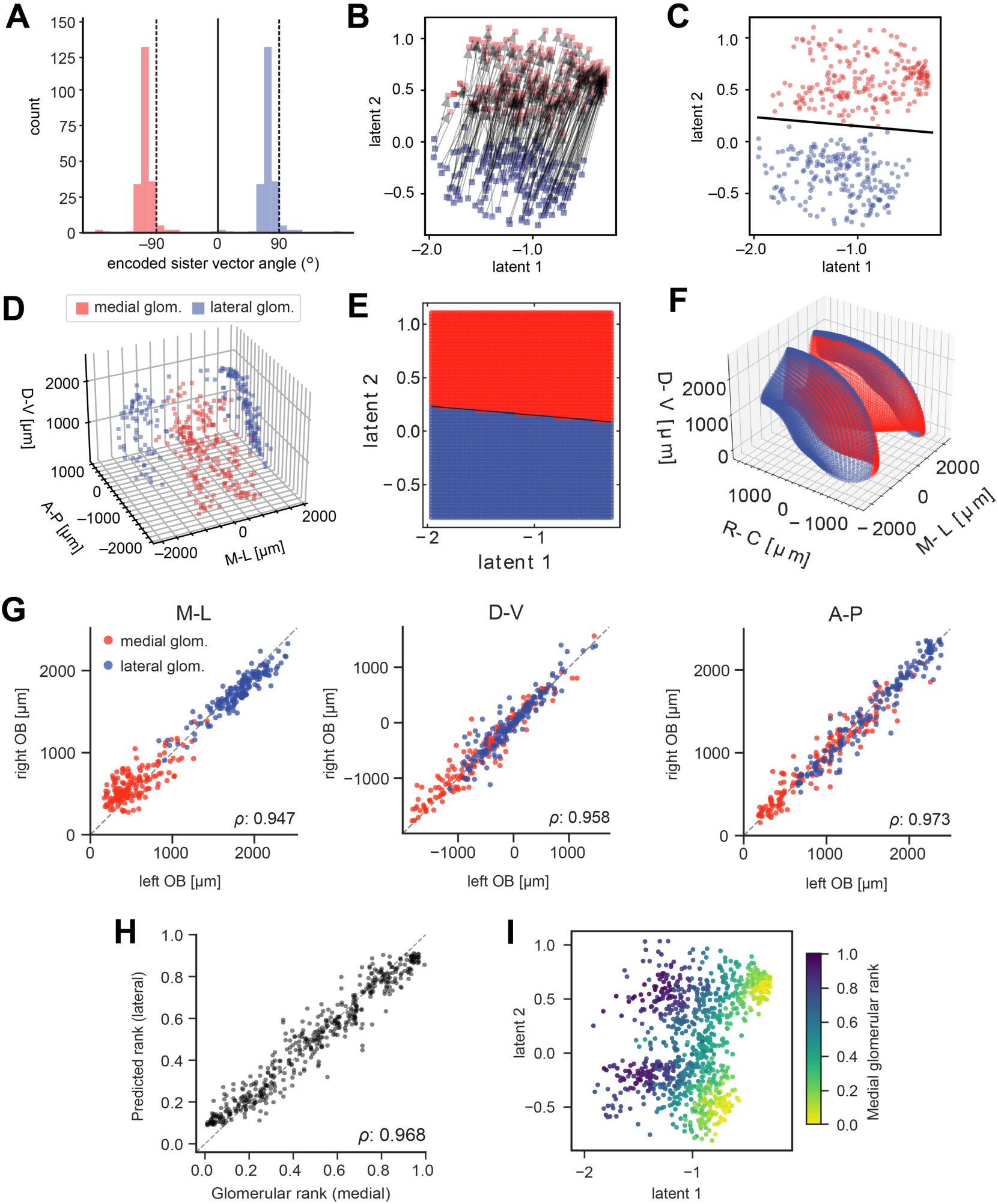
Symmetry of the glomerular map revealed by positions of sister glomeruli. **(A)** In the latent space encoding produced by the dual autoencoder, the relative vector angles of sister glomeruli are well separated into two clusters. **(B)** Curated medial (red) glomeruli share the same vector directions in the latent representation and are spatially separated from lateral (blue) glomeruli. **(C)** Linear separation of the medial (red) and lateral (blue) glomerular domains in the latent map defines the intra-hemispheric axis of symmetry (black line). **(D)** Medial and lateral sister glomeruli of the curated input glomerular dataset in 3D across left and right OB. **(E)** The linear discriminant allows assigning any location in the glomerular manifold to either of the two glomerular domains. **(F)** Visualization of the entire 3D glomerular manifold with spots colored by domain identity in latent space. **(G)** Linear correlation between glomerular maps in the left and right OB across the medial-lateral (M-L), dorsal-ventral (D-V), and anterior-posterior (A-P) axes. Each dot represents a single glomerulus and is colored by glomerular domain. ρ = Pearson’s correlation coefficient. **(H)** Linear correlation between the rank of glomeruli in the medial glomerular domain and predicted lateral glomerular ranks. For details on glomerular ranking see Methods. **(I)** Positions of the 968 lateral and medial glomeruli in latent space colored by medial glomerular rank.

We next analyzed the relative positions of sister glomeruli. This analysis revealed that sister glomeruli expressing the same OR are clearly separable along a medial-to-lateral axis within each hemisphere (**Fig. 4G**). To characterize the organization of individual glomeruli along this axis, we ranked the positions of glomeruli within the medial domain (see Methods). Glomerular ranks were well-preserved when compared along the lateral domain. Support vector machines trained on lateral glomerular positions could accurately predict the medial glomerular ranks of lateral glomeruli (**Fig. 4H and I**). Together, our data provide a detailed, quantitative description of glomerular identity and position, revealing a glomerular map that is mirror-symmetric between hemispheres and duplicated with a remarkably high fidelity within each OB.

### The molecular organization of the glomerular map

Prior work has implicated OR sequence similarity and chromosomal location, as well as axon guidance gene expression in the OE, in determining the broad structure of the glomerular map, such as its organization into dorsal and ventral domains^13,15,30–36^. However, how these factors contribute to the positioning of individual glomeruli across the entire glomerular map is unknown. Therefore, we next sought to determine the extent to which OR identity and OSN gene expression predict glomerular position (**Fig. 5A**). Canonical ORs can be divided into two classes, with approximately 130 class I and 950 class II ORs^37,38^. Consistent with prior work^39,40^, glomeruli expressing class I ORs were almost exclusively found in the dorsal-most part of the OB, while glomeruli expressing class II ORs were found throughout the rest of the OB (**Fig. 5B**). For class II ORs, we observed a moderate but significant correlation between their phylogenetic similarity and glomerular positions, with more similar pairs of ORs located closer in space (**Fig. 5C and D**). Furthermore, we found that the pairwise glomerular distance distributions were smaller on average for OR pairs within the same chromosomal cluster (**Fig. S9A and B**). Finally, chromosomal clusters with ORs that were phylogenetically more similar had, on average, closer pairs of glomeruli (**Fig. 5D-F**). For example, a population of glomeruli expressing a cluster of phylogenetically similar ORs located on chromosome 10 was restricted to the anterior portion of the OB; this cluster corresponds to the recently identified subtype of OSNs that express the transmembrane protein CD36^41^ (**Fig. 5F**). These results suggest a significant correlation between OR sequence similarity and genomic proximity with glomerular position for glomeruli/receptors within the class II domain.

**Figure 5:**
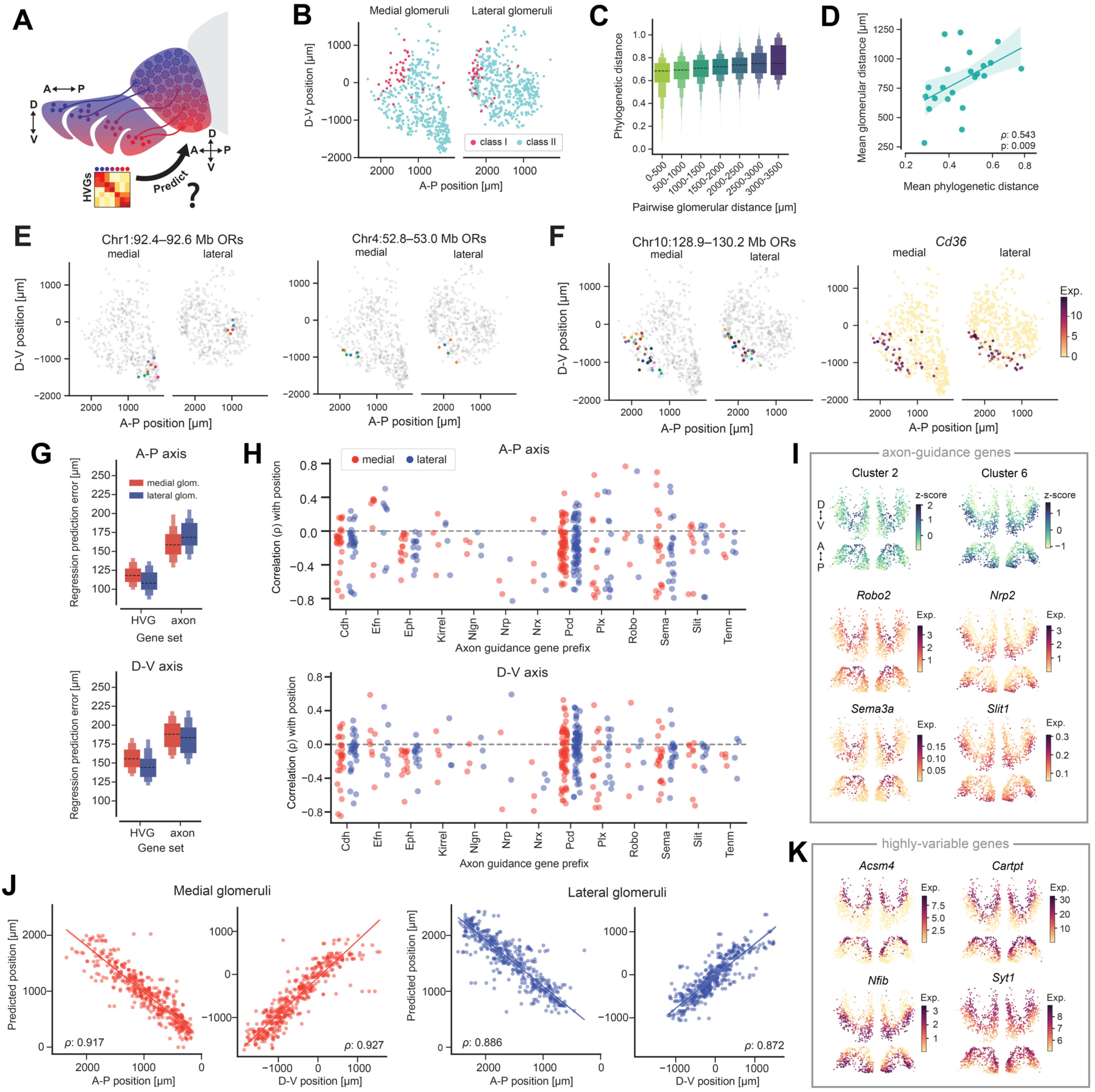
OR gene identity and non-OR genes explain glomerular position. (**A**) Schematic representation of the objective to determine how the unique transcriptomes of OSNs in the OE, and their expression of highly variable genes (HVGs), predict glomerular position in the OB. (**B**) Positions of glomeruli expressing class I (magenta) and class II (cyan) OR genes. (**C**) Letter value boxplot of phylogenetic distance of ORs expressed in class II glomeruli compared to pairwise glomerular distance (in µm), revealing that glomeruli expressing more phylogenetically similar class II ORs are located closer to each other in the OB. (**D**) Correlation between mean phylogenetic distance and glomerular distance across chromosomal clusters (blue points). (**E**) Spatial position of glomeruli expressing ORs in chromosome 1:92.4-92.6 Mb (left plot) and chromosome 4:52.8-53.0 Mb (right plot), showing that glomeruli expressing these ORs are located close together in the OB. (**F**) Spatial position of glomeruli expressing ORs in chromosome 10:128.9-130.2 Mb (left plot) and expression of *Cd36* (right plot) show a high correlation. (**G**) Letter value boxplot of regression prediction error from elastic net regularized linear regression models trained on either a set of axon guidance genes or HVGs (highly variable genes) to predict glomerular positions. No significant difference exists between regression errors in predicting positions of glomeruli in the medial or lateral domains, but lower regression errors for HVGs indicate a higher accuracy in predicting glomerular positions based on this gene set. (**H**) Dotplot of gene correlation for medial glomeruli (red) and lateral glomeruli (blue) for individual axon guidance gene families. Top plot shows correlations along the D-V axis and bottom plot shows correlations along the A-P axis. (**I**) **Top:** Hierarchical clustering of axon guidance gene expression patterns revealed clusters that map onto the A-P (Cluster 2) and D-V (Cluster 6) axes of the OB. **Bottom:** Expression of individual axon guidance genes. In each subpanel, top plot shows Z-projection and bottom plot shows Y-projection of the OB (**J**) Correlations of predicted positions of glomeruli using a regression model trained on the HVG set. The model performed slightly better at predicting positions of medial glomeruli (red) across both the D-V and A-P axes (ρ=0.927 and 0.917, respectively) compared to lateral glomeruli (blue, ρ=0.872 and 0.886, respectively). (**K**) Expression of individual highly variable genes. Top plots show Z-projection and bottom plots show Y-projection of the OB.

We next examined the relationship between the molecular organization of the glomerular map and gene expression programs in the OE. Recent work using single-cell RNA-sequencing (scRNA-seq) revealed a surprising link between OR identity and OR-specific gene expression programs^17,33,34^. Therefore, we asked the extent to which these unique OSN transcriptomes are predictive of glomerular position. We first selected 180 well-characterized axon guidance and cell adhesion molecules (**Table S1**) and assessed how well the expression of these genes, measured via scRNA-seq of OSNs, could predict the location of glomeruli, measured via spatial transcriptomics. We observed that the expression of these axon guidance genes strongly correlated with glomerular positions (**Fig. 5G-K**). Furthermore, individual gene families had similar correlations with positions from the lateral and medial glomeruli, suggesting that such genes may influence the observed symmetry in the glomerular maps across the medial and lateral hemispheres (**Fig. 5H**). Elastic net regularized linear regression models trained on these axon guidance genes accurately predicted glomerular positions for each of the anterior-posterior and dorsal-ventral axes, with an average regression prediction error of 170.9 µm for lateral glomeruli and 160.3 µm for medial glomeruli along the A-P axis, and 183.3 µm for lateral glomeruli and 189.2 µm for medial glomeruli along the D-V axis (**Fig. 5G, Fig. S9C and D**). Hierarchical clustering of the expression patterns of these genes revealed sets of genes with distinct patterns of correlations with glomerular positions, two of which mapped onto the anterior-posterior and dorsal-ventral axes (**Fig. 5I**). These clusters contained axon guidance genes that have previously been shown to regulate glomerular positions, such as *Robo2*, *Nrp2 and Slit1*^42–44^, as well as genes whose functions in glomerular map formation remain uncharacterized (**Fig. 5I**, **Fig. S10A**).

Finally, we asked whether the transcriptional variation across OSNs is reflected in the glomerular map. We recently characterized ∼1,300 highly-variable genes (HVGs) that could accurately distinguish between OSNs expressing different ORs^34^. We therefore tested the extent to which the expression of these HVGs could predict glomerular positions. We observed that HVG expression correlated with glomerular positions and that elastic net models trained on their expression across OSN subtypes could accurately predict the glomerular positions of their respective ORs, with an average regression prediction error of 110.5 µm for lateral glomeruli and 118.8 µm for medial glomeruli along the A-P axis, and 146.7 µm for lateral glomeruli and 157.4 µm for medial glomeruli along the D-V axis (**Fig. 5G and J, Fig. S9C**). Examination of HVGs predictive of glomerular position identified both known and novel genes. For example, the family of NFI transcription factors predicted the positions of ventral glomeruli, whereas the expression of genes like *Acsm4* and *Cartpt* was enriched in OSN subtypes that project to the dorsal part of the OB^45,46^ (**Fig. 5I**). Additionally, genes downstream of OR signaling, like phosphodiesterase enzymes and *Pkia,* were also predictive of glomerular positions (**Fig. S10B**). This observation was especially robust along the anterior-posterior axis, consistent with prior work demonstrating how glomerular positions along this axis are influenced by ligand-independent OR activity^47^ (**Fig. S10B**). Finally, we observed that the expression of synaptic genes like *Syt1* varied as a function of projection patterns along the dorsal-ventral axis, suggesting ways in which OSN function might vary across the OB (**Fig. 5K**).

In summary, by integrating transcriptomics datasets from the OE and OB, we reveal that transcriptional differences across OSN subtypes reflect their connectivity with distinct glomerular domains, are predictive of the stereotyped glomerular positions along each OB axis, and may correspond to differences in OSN function and signaling along these axes.

### Mitral cell projection neurons and deep-layer granule cells exhibit molecular structure along alternative axes

Finally, we asked whether the symmetry of the glomerular map is reflected in molecularly distinct domains of deeper layer mitral and granule cells (**Fig. 6A**). To restrict our analysis to defined cell types and layers, we isolated capture spots from the Leiden Clusters (**Fig. 1D and E**, **Fig. 6B**). From these clusters, we selected those representing mitral cells (Leiden Cluster 6), superficial granule cells (Leiden Clusters 8 and 9), and deep granule cells (Leiden Cluster 1) based on their alignment with morphology (**Fig. 6B and C**). We projected the axis of glomerular symmetry onto the deeper layers, and we performed differential gene expression analysis across this axis, as well as across alternative axes (see Methods, **Fig. 6A and D**).

**Figure 6:**
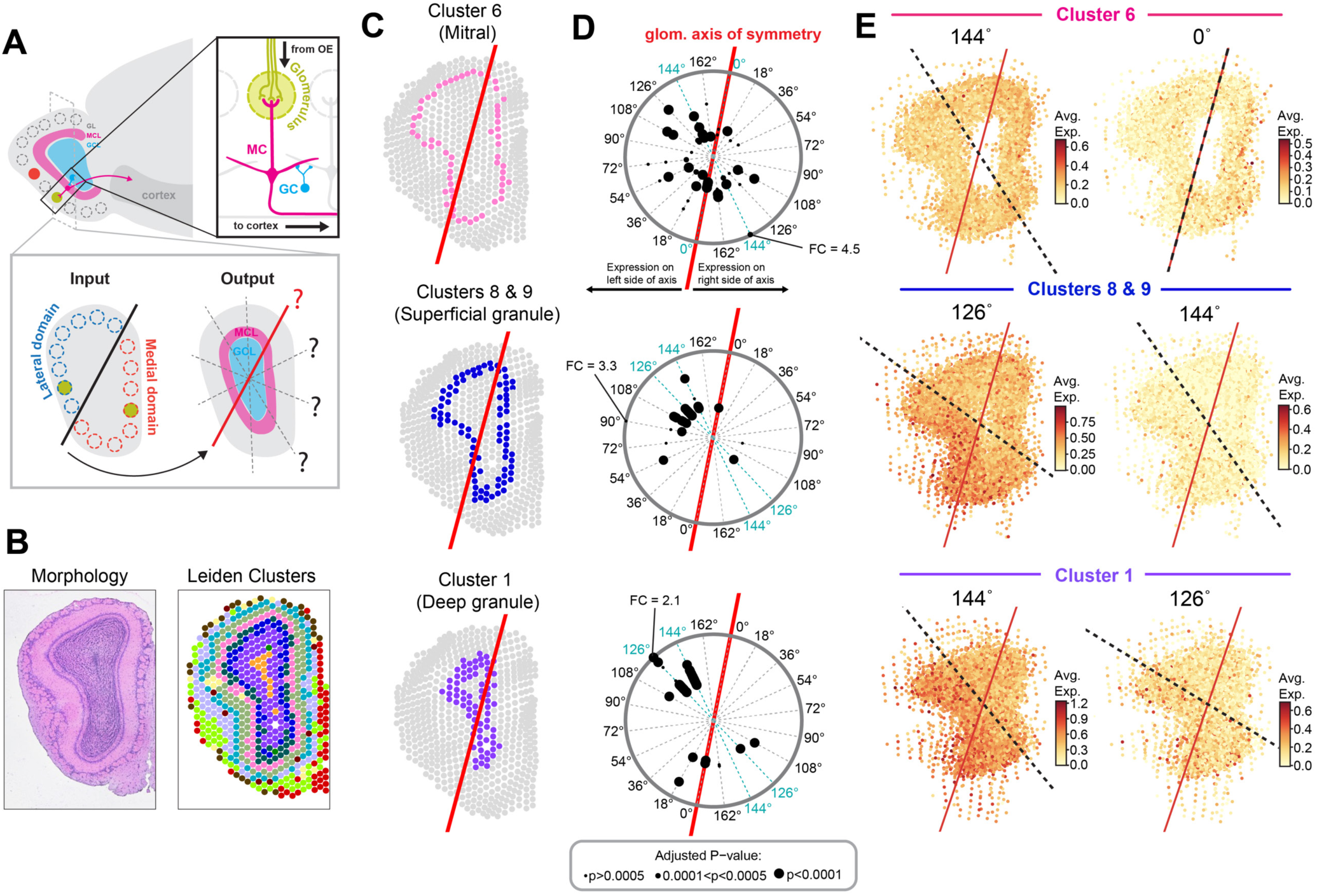
Gene expression in mitral and granule cell layers is distributed across axes alternative to the glomerular axis of symmetry. **(A) Top**: Schematic of output circuit from OB to cortex where microcircuits containing mitral cells (MC, pink) send projections to cortex, whose activity is shaped by inhibitory interneurons including granule cells (GC, blue). **Bottom**: Schematic of the research objective to determine whether the glomerular axis of symmetry is reflected in deep layer cells through differential gene expression, or whether there are alternative axes that structure these layers. **(B)** Representative image of morphology (left) and capture spots in one section of the left OB with Leiden clusters (right) **(C)** Capture spots in a single section colored by isolated Leiden cluster. Clusters isolated for analysis represent the mitral cell layer (Cluster 6), superficial granule cell layer (Clusters 8 & 9), and deep granule cell layer (Cluster 1). The glomerular axis of symmetry is overlaid in red. **(D)** Polar plots showing the axis angle for the top 50 genes that maximize differential gene expression across the axis (see Methods). Red line is the glomerular axis (0°). Size of points represents adjusted p-value and radial location represents the log_2_ fold change. Genes with the highest absolute log2 fold change are indicated at the edge of the polar plots and fold change (FC) is provided. Angular location indicates the axis angle of maximal symmetry. Points located on the right half of the polar plot indicate genes upregulated on the right side of the axis, and points on the left half of the polar plot are genes upregulated on the left side of the axis. **(E)** Expression signatures of the two peak axis angles within the isolated capture spots for each Leiden cluster flattened in 2D. Normalized expression is averaged across all genes within the gene signature.

Our analysis revealed significant differential gene expression along multiple different axes. Both superficial and deep granule cell layers exhibited preferential alignment of gene expression along two axes alternative to the glomerular axis, at angles 126° and 144° from the glomerular axis (Red line, angle 0°) **(Fig. 6D and E, Fig. S11**). However, gene expression patterns in the mitral cell layer were much more diverse, with genes aligned along multiple different axes (**Fig. 6D and E, Fig. S11, Table S2**). In all three layers, gene expression did not preferentially align with the glomerular axis of symmetry (**Fig. 6D)**. These data suggest that the symmetry defined by OR identity in the glomerular layer is reorganized, rather than mirrored, in the mitral and granule cell layers.

## DISCUSSION

We here provide a detailed 3D map of the molecular structure of the mouse OB and determine the precise positions of hundreds of glomeruli. The spatial arrangement of molecularly distinct glomeruli defined two axes of symmetry. The first was the midline, dividing mirror-symmetric glomerular maps in the left and right OB, while the second was within each OB, at the boundary between the medial and lateral duplications of the map. The organization of mitral and granule cells did not mirror this glomerular layer symmetry. By integrating spatial transcriptomics and single-cell RNA sequencing data, we found that gene expression programs in the OE could determine glomerular positions with near-glomerular resolution.

### Symmetry of the glomerular map

In the OB, glomeruli and their connected microcircuits are functional units that route odor information from primary sensory neurons in the OE to the olfactory cortex. An intriguing feature of the glomerular map is its apparent mirror symmetry across hemispheres and within each OB. Based on positional information from hundreds of glomeruli and a dual autoencoder model to account for the complex geometry of the glomerular layer, we identify a curved plane as the axis of this intrahemispheric symmetry, based on the relative positions of sister glomeruli expressing the same OR along the curved plane. This analysis revealed that the locations of sister glomeruli were aligned with precision along a medial-to-lateral axis of symmetry, such that the ranks of lateral glomeruli can be accurately predicted from the ranks of medial glomeruli.

Prior work suggests that mirror symmetry in the glomerular map may be critical for olfactory navigation^48–52^. Animals utilize olfactory stereo-sampling to sample dynamic odor plumes, and OSNs from the medial (septum) and lateral (turbinate) olfactory epithelium project to the medial and lateral OB^13^. Thus, differences in odorant-receptor interactions across the olfactory epithelium may result in the differential activation of the lateral and medial glomerular domains of the OB. The axis of intra-hemispheric symmetry we define will allow for future research to elucidate the functional relevance of mirror-symmetric glomerular maps. More generally, the glomerular map provides a unique experimental model for determining the molecular mechanisms that establish or disrupt symmetry in the brain, for example by manipulating the expression of symmetry-breaking genes^53–55^.

What is the relationship between the glomerular map and the organization of neurons in the deeper layers of the OB? Each glomerular column contains a set of mitral and tufted cell projection neurons that route information to downstream cortical areas, and molecularly distinct mitral and tufted cell subtypes preferentially project to different downstream cortical regions^25,56,57^. Moreover, mitral and tufted cell activity is modulated by local inhibitory neurons, including periglomerular cells and deep-layer granule cells^58–60^. While our analysis reveals spatial structure of gene expression in the mitral and granule cell layers, our results suggest that the axis of glomerular symmetry is reorganized, rather than mirrored, in the molecular patterns of output neurons. This reorganization may provide a mechanism to decorrelate the duplicated glomerular input channels.

### The molecular structure of the glomerular map

A range of diverse molecular mechanisms has been proposed to control glomerular positioning, including OR identity and chromosomal location, intrinsic signaling, and graded expression of axon guidance genes during development^33,61–63^. However, prior studies perturbing the expression of individual ORs or axon guidance genes resulted in only partial disruptions to the glomerular map^32,35,63–68^. Thus, precise glomerular positioning likely relies on multiple complementary mechanisms for axon guidance.

By utilizing sequencing-based spatial transcriptomics, we gain access to the entire OB transcriptome, allowing us to comprehensively assess gene expression programs along axes of glomerular symmetry. We identified a modest but significant correlation between glomerular position and phylogenetic OR similarity, consistent with a model that comprehensive gene expression programs beyond OR identity determine glomerular position^17,18,35^. We next assessed whether the unique transcriptomic identity of OSNs contributes to glomerular positioning. We found that the expression of a set of characterized axon guidance genes and cell adhesion molecules predicted the position of glomeruli with an accuracy of ∼200 µm. This prediction accuracy was further improved to 120-150 µm, near glomerular accuracy, when including a set of highly variable genes (HVGs) expressed in OSNs in the OE^34^. Together, these results suggest that gene expression programs in the OE, which provides sensory input to the OB, are strongly correlated with the molecular organization of the glomerular map.

In summary, the ability to map the organization of glomeruli, the structural and functional units of information processing in the olfactory bulb, has revealed the striking precision with which molecular information determines the spatial organization of highly symmetric glomerular maps. By integrating this structural information with functional approaches, future experiments will allow us to unravel the molecular-anatomical basis for information processing in the olfactory system.

## AUTHOR CONTRIBUTIONS

Conceptualization: NK, MK, ATS, AF; methodology: NK, MK, DHB, DB, CT, YM; investigation: NK, MK, DHB, DB, CT, YM; visualization: NK, MK, DHB, CT; funding acquisition: NK, ATS, AF; project administration: AF, ATS; supervision: NK, MK, YM, ATS, AF; writing - original draft: NK, MK, DHB, ATS, AF; writing - review & editing: NK, MK, DHB, CT, DB, YM, ATS, AF

## ACKNOWLEDGEMENTS

We thank Joakim Lundeberg, Kim Thrane, and Annelie Mollbrink from the Department of Gene Technology, KTH Royal Institute of Technology in Stockholm, Sweden for support in optimizing the spatial transcriptomics protocol to capture OR transcripts in OB. We thank Lindsay Scarpitta and Pieter Faber from the Genomics Facility at the University of Chicago for sequencing services, and Christoph Schorl from the Genomics Facility at Brown University for quality control of cDNA libraries. We thank Nicole Eckart from 10x Genomics for excellent technical support, Sina Tootoonian for advice on modeling, and Carles Bosch from the Crick Institute, as well as Norman Rzepka and Florian Meinel from Scalable Minds for their help with tissue alignment. We thank David Berson, Tom Bozza, Kevin Franks, and Anthony Crown for critical comments on the manuscript, and we thank Gilad Barnea for providing antibodies and for critical comments on the manuscript.

## RESOURCE AVAILABILITY

### Lead contact

Requests for further information and resources should be directed to and will be fulfilled by the lead contact, Alexander Fleischmann.

### Materials availability

This study did not generate new unique reagents.

### Data and code availability

Raw and processed spatial transcriptomics sequencing data have been deposited in Gene Expression Omnibus (GEO) and will be publicly available upon publication.

All original code used for this paper will be publicly available upon publication.

### Additional information

Any additional information required to reanalyze the data reported in this paper is available from the lead contact upon request.

## Supplemental information

**Table S1**: Axis correlation scores and f-scores for 180 axon guidance genes, related to Figure 5.

**Table S2**: Top 50 differentially expressed genes by Log_2_FC per Leiden cluster, with axis angle, related to Figure 6.

## METHODS

### Experimental model and subject details

One male WT C57Bl/6 mouse (7-weeks old) was used in this study and obtained from Jackson Laboratories. All animal protocols were approved by the Brown University’s Institutional Animal Care and Use Committee (protocol number: 21-03-0004) followed by the guidelines provided by the National Institutes of Health.

### Spatial transcriptomics of mouse olfactory bulb

#### Tissue dissection and sectioning

Whole olfactory bulb tissue was dissected, embedded in OCT and frozen in a bath of 2-methylbutane surrounded by powdered dry ice. 200 consecutive 10µm sections of tissue were sectioned on a Leica CM3050 S cryostat, mounted onto Visium Spatial Gene Expression slides (10X Genomics) and maintained at −80°C until processing. Every fifth 10 µm cryosection was mounted onto a standard Superfrost microscope slide (Fisher Scientific) and saved at −80°C to use for OR expression validation using immunohistochemistry with OR antibodies against Olfr6 (M50) and Olfr1507 (MOR28). A small set of tissue sections (17) were not mounted on Visium slides or standard microscope slides due to damage. These sections were accounted for as 10 µm ‘gaps’ in the alignment of the 3D OB dataset.

#### Fixation, staining, and imaging of tissue sections

The 200 sections were processed according to the Visium Spatial Gene Expression protocol (10x Genomics) with slight modifications. Prior to the experiment, a preliminary tissue optimization experiment was performed to determine the minimum tissue permeabilization time to yield high capture of OR mRNA from delicate OSN axon termini. Tissue sections from 4 Visium slides (16 sections total) were processed each day. Slides were removed from −80°C, tissue sections were dried by incubating on a thermal cycler at 37°C for 1 minute, and were then fixed in methanol for 30 minutes in −20°C. The sections were then stained with Hematoxylin and Eosin according to the 10X Visium protocol and imaged using a Nikon Eclipse Ti2 inverted microscope to obtain reference histology images for each tissue section.

Tissue sections were then permeabilized for 6 minutes according to the preliminary optimization experiment, allowing for mRNA to diffuse and be captured by barcoded mRNA capture probes printed on the Visium slides. The captured mRNA transcripts were then reverse-transcribed, and second-strand cDNA was generated. Then, qPCR was performed to determine C*_t_* values for each sample, and PCR amplification was performed on the samples.

Barcoded cDNA libraries of the 200 tissue sections were generated using the Spatial Gene Expression reagents (10X Genomics) according to the manufacturer’s protocols. Libraries were pooled and sequenced on an Illumina NovaSeq 6000 instrument using the following run variables: 28 cycles (Read 1); 10 cycles (i7 index); 10 cycles (i5 index); 120 cycles (Read 2). The samples were initially sequenced to achieve an average depth of 50,000 read pairs per capture spot, then sequenced again to achieve a final average depth of 200,000 read pairs per capture spot.

#### Data pre-processing

Data were demultiplexed and processed using SpaceRanger v1.1.0. For quality control checks of non-OR gene expression in the OB, reads were aligned to the mouse genome (mm9). The reads were pseudoaligned to the mouse transcriptome using the alevin-fry pipeline to account for the highly homologous sequences of OR genes^69^.

#### Visualization of gene expression in individual sections

Analysis of spatial transcriptomics data in individual tissue sections was performed in R using Seurat V4. Expression data was normalized using SCTransform where indicated and expression of individual genes (layer marker genes and OR genes) was overlaid against H&E histology images. For the identification of glomerulus positions, raw (non-normalized) and non-filtered data of OR gene expression were used due to the highly variable and sparse capture of OR genes.

#### Integration of capture spots and deconvolution with scRNA-seq OB dataset

Gene expression from 190,445 capture spots across the 200 spatial transcriptomics datasets was integrated using the scVI analysis pipeline from scvi-tools. The clustering of capture spots was performed using the Leiden algorithm, producing 16 clusters.

Deconvolution and visualization of the integrated spatial transcriptomics dataset was performed using Deconvolution of Spatial Transcriptomics profiles using Variational Inference (DestVI) from scvi-tools. The single-cell model (scLVM) was trained using an integrated OB scRNA-seq dataset^25,70^, and then the spatial model (stLVM) was trained to perform the deconvolution.

#### Immunohistochemistry to validate OR transcript capture

10µm tissue sections saved in −80°C for validation were first dried for 1 min at 37°C and fixed for 7 min in 2% PFA. Sections were washed in PBS three times for 10 min, permeabilized in PBS and 0.1% Triton X-100 for 30 min, then blocked in PBS, 0.1% Triton X-100 and 5% heat-inactivated horse serum at room temperature for 30 min. Sections were then incubated overnight at 4°C with primary antibodies against M50 (guinea pig, 1/1000, Gilad Barnea) and MOR28 (rabbit, 1/1000, Gilad Barnea). Sections were then washed with PBS and 0.1% Triton X-100 three times for 10 min, blocked in PBS, 0.1% Triton X-100 and 5% heat-inactivated horse serum at room temperature for 30 min, then incubated with secondary antibodies, anti-guinea pig conjugated to 488 and anti-rabbit conjugated to Cy3 (1/1000, Donkey IgG (H+L), Jackson ImmunoResearch) for 2 hours at room temperature in the dark. Sections were rinsed three times with PBS and 0.1% Triton X-100 and sealed with Fluorescent Vectashield Mounting Medium (Vector, H-1900), and imaged at 20X using a Nikon A1R0HD confocal microscope.

### Three-dimensional alignment of morphology and gene expression

#### Manual annotation and alignment of morphological sections

Manual annotation was necessary to account for stretching, folding, and damage of tissue sections during mounting. Initial alignment was performed by first conducting coarse alignment through grey-level thresholding to identify tissue areas, followed by cropping a square region centered on the centroid of the segmented object. Misaligned tissue sections were manually corrected by flipping them horizontally or vertically as needed. Subsequently, a custom FIJI script^71^ was employed to align the image stack using BUnwarpJ^72^, an elastic registration tool. Finally, the transformations generated by BUnwarpJ were applied to the corresponding spot coordinates.

Further refinement was performed via manual annotation of glomeruli using WebKnossos and the data annotation service Mindy^73^. First, based on the distribution of nuclei in the reference histology images, the centers of 2,550 glomeruli were annotated in individual sections across the 200 tissue sections. Then, glomeruli were annotated by identifying the centers of the same glomeruli across consecutive sections. The resulting glomerular annotations were then used to refine the morphological alignment using AI computer vision with Scalable Minds, by performing fine-scale spatial transformations of the tissue sections.

#### Alignment of gene expression data in 3D

The gene expression data was aligned in 3D according to the morphological alignment, by transforming the capture spot position of the gene expression data to the aligned morphological images. The z-position for the 3D dataset was approximated based on the thickness of tissue sections (10µm).

#### Categorization of variable OR expression

Genes with total expression values below 5 were assigned to the “none” cluster. For the remaining (expressing) genes, we computed three metrics: clusteredness, total expression, and number of nonzero counts, which we transformed using a hyperbolic arcsine function and normalized to the [0,1] range. We then applied agglomerative clustering (using Euclidean distance and Ward linkage) to partition the expressing genes into four groups. To assign biological meaning to these clusters, we ranked the clusters based on their mean total expression and clusteredness. The cluster with the highest difference between ranked expression and ranked clusteredness (indicating high expression but low spatial clustering) was labeled “diffuse”. Among the remaining clusters, the one with the highest ranked expression was labeled “high,” followed by “medium” and “low” for the clusters with progressively lower ranked expression. The clusters defined here were only used for the visualization of scope expression patterns among the 1135 genes and were not used in subsequent analysis.

### Computational modeling to quantify glomerular position

#### Probabilistic Glomerulus Position (PGP) Model

To identify the precise locations of olfactory glomeruli from the spatial transcriptomics data, we developed a model that predicts the expected gene counts at each spot based on the positions and radii of putative glomeruli. The model assumes that the count value at a spot is proportional to the overlap between the spot’s surface (modelled as a disk) and the glomerulus (modelled as a sphere). We define the overlap function 𝑂(𝑅_*g*_, Δ_*x*_, Δ_*y*_, Δ_*z*_) that calculates the proportion of the spot area occupied by a glomerulus with radius 𝑅_*g*_, Δ_*x*_, Δ_*y*_ and Δ_*z*_ representing the distances between the spot and glomerulus centres in the x, y, and z directions, respectively. The overlap function is given by:

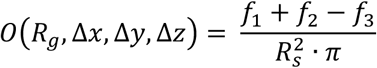

where 𝑅*_S_* is the spot radius, and 𝑓_1_, 𝑓_2_ and 𝑓_3_ are calculated as follows:

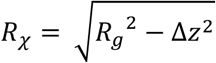

(The radius of the circular cross-section of the glomerulus in the plane of the spot.)

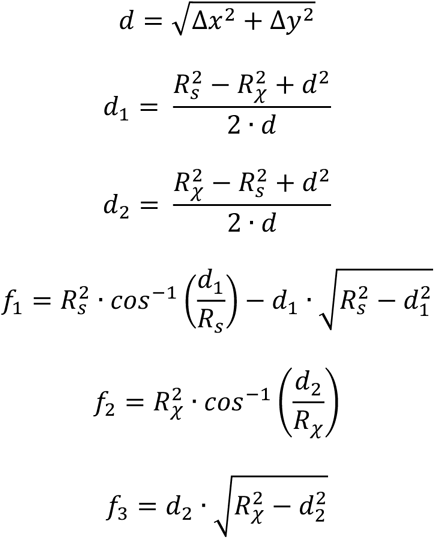

The count value 𝐶*_n_* at spot 𝑛 is pulled from a normal distribution (allowing for non-integer count values) with a mean determined by the sum of overlaps with all 𝐾 glomeruli, their respective expression strengths 𝑆, and the binary variable 𝑉 ∈ {0,1} representing the validity of the glomerulus (whether it should be considered), calculated from a glomerulus validity probability value 𝑉 = 𝐵𝑒𝑟𝑛𝑜𝑢𝑙𝑙𝑖(𝑝)).

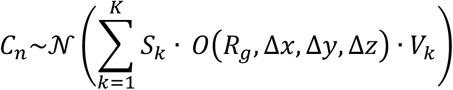

The above functions were implemented in Julia (version 1.9.2). Probabilistic modeling and inference were performed using Turing.jl (version 0.28.2). The model parameters, including glomerulus centers (𝑥, 𝑦, 𝑧) and validity probabilities 𝑝, are inferred using a particle Markov chain Monte Carlo (MCMC) algorithm.

Prior putative glomerular locations were initialized for optimal computational efficiency based on the raw spot count values by a lenient heuristic method. Initial peaks in the gene expression data were identified by comparing the expression values to a detection threshold (20% of the mean expression value). To determine the strength value of each glomerulus for the prior distribution, the overlap function 𝑂(𝑅_*g*_, Δ_*x*_, Δ_*y*_, Δ_*z*_) was used (for the heuristic function) to update the count value of each spot based on their distance from neighboring spots and their count values. Peaks with expression levels below 0.2 were discarded, and all other peaks were clustered based on their Euclidean distance. The median coordinates of peak clusters within 100 µm were used as the expected value of glomerular centers, and the maximal peak expression value of the spots within the cluster was used as the estimated strength value. Priors were defined for the center coordinates as normal distributions with a 200 µm standard deviation, and the mean was defined by the heuristic described above. Log-normal priors were used for the radii (50 µm expected value) and strength (as defined by the heuristic function). For the glomerulus probability, a Kumaraswamy distribution was used (instead of a Beta distribution to improve computational efficiency), with a variance of 0.1 and expected value based on the relative strength of the glomerulus compared to the maximal count value in the gene’s dataset 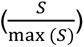.

The probabilistic model was sampled for 10,000 iterations for each gene with a particle Gibbs sampler using 30 particles in parallel.

#### Human Expert Curation of High-Confidence Glomeruli

The curated dataset was generated by simultaneously displaying the entire raw dataset for each gene in an interactive navigable 3D plot together with the proposed glomeruli and their overlap. Each putative glomerulus was confirmed if the glomerular signal was clearly visible at the center of the putative glomerulus and not associated with afferent axonal fibers or background.

#### Symmetrization of Left and Right Hemi-Bulbs

First, manually curated divider points separating the left and right bulbs were annotated in WebKnossos^74^. A curved divider surface was fitted to the divider points using a dense mesh grid (L-BFGS-B minimization algorithm). All glomeruli in the output of the PGP model were assigned *right* or *left* laterality according to their position relative to the divider surface. For symmetrization, a cost function was defined to measure the dissimilarity between the positions of glomeruli in the left and right bulbs after applying a transformation. Deviations from symmetry were minimized using the L-BFGS-B algorithm to find the optimal symmetrization vector that aligns the glomeruli in the left and right bulbs. The glomeruli positions in the right olfactory bulb were transformed using the estimated symmetrization vector. The transformed right glomeruli positions were concatenated with the original left ones to create a symmetrized dataset.

#### Double Autoencoder (AE) Model for Sister Glomerulus Positions

To model the relative positions of sister glomeruli in the medial and lateral domains of the olfactory bulb, we developed a dual autoencoder architecture consisting of a two-layer encoder network and two decoder networks. The model aims to:

1. Map the 3D positions of glomeruli onto a 2D latent space (embedding) from which it can reliably reconstruct the original positions. 2) Predict the position of the sister glomerulus for each glomerulus. 3) Ensure that the mapping of glomeruli and sister glomeruli are similar in the latent space.

The dual autoencoder model was designed with minimal trainable parameters to allow generalization on the curated glomerulus dataset. It consists of the following components:

<u>Encoder:</u> A two-layer (21 and 7 neurons) dense neural network that maps the 3D glomeruli positions to a lower-dimensional latent space. The encoder layers use sigmoid activation function and L2 regularization.

<u>Decoder:</u> A two-layer dense neural network that reconstructs the 3D glomeruli positions from the latent space representation. The decoder layers are tied to the encoder layers, sharing the same architecture and weights.

Sister Glomerulus Predictor: A single dense layer with linear activation that predicts the latent space representation of the sister glomerulus given the latent space representation of a glomerulus.

<u>Sister Glomerulus Decoder:</u> A decoder network that reconstructs the 3D position of the sister glomerulus from its predicted latent space representation. It is identical to the main Decoder and shares the same weights.

The dual autoencoder model is trained using a compound loss function combining:

Reconstruction Loss: Mean squared error (MSE) between the input 3D glomeruli positions and their reconstructed positions from the autoencoder.

Sister Glomerulus Prediction Loss: MSE between the predicted 3D positions of sister glomeruli and their true positions.

Encoding Similarity Loss: MSE between the predicted latent space representation of the sister glomerulus (from the Sister Glomerulus Predictor) and its encoding from tissue coordinates (from the Encoder).

The resulting latent space representation can reveal clusters or patterns related to the medial and lateral domains of the olfactory bulb, allowing for unsupervised assignment of medial/lateral labels to each glomerulus based on their relative positions to their sisters. The trained dual autoencoder model can be used for various tasks, such as: Encoding glomeruli positions to the latent space, decoding latent space representations to reconstruct glomeruli positions, predicting the positions of sister glomeruli given a glomerulus position, and inferring the medial/lateral labels of glomeruli based on their relative latent space representations.

#### Confirmation of Glomeruli using PGP and AE Model Predictions

To confirm the validity of the glomeruli identified by the PGP model and identify the most likely set of glomeruli for each gene, we utilized the predictions from the AE model. The confirmation process was performed using the following steps:

Each gene’s dataset was separately analyzed. Mirror positions of the glomeruli were calculated by negating the x-coordinate of the left glomeruli and assigning it to the right glomeruli after symmetrization. The Euclidean distances between the mirror positions of the left and right glomeruli were calculated. These distances represent the mirror errors, indicating how well the putative glomeruli align with their mirror counterparts. The dual autoencoder model was used to predict the positions of the sister glomeruli for each putative glomerulus. The Euclidean distances between the predicted putative sister positions and the actual glomeruli positions were calculated. These distances represent the sister errors, indicating how well glomeruli aligned with the predicted other glomeruli on the same side.

Glomeruli were considered valid if they satisfied the following criteria:

1. Either the sister error or the mirror error was less than 250 microns. 2) The probability of the glomerulus, as predicted by the PGP model was greater than 0.25.

For each gene, combinations of valid glomeruli were exhaustively searched to find the combination with the minimum total error, considering both sister errors and mirror errors. The search started with combinations of 4 glomeruli and decreased to 2 glomeruli if a satisfactory combination was not found.

### Ranking sister glomeruli

Medial geometrically-confirmed glomeruli were ranked along a single axis using a nearest-neighbor approach. 3-D glomerular positions were scaled using scikit-learn’s RobustScaler, and a k-nearest neighbor graph was constructed using k=10 neighbors. To obtain a stable ranking of glomeruli, a vector the size of the number of glomeruli was constructed with all zeros except for the most medial glomerulus. This vector was then multiplied by the kNN graph (weighting each cell by 0.1 and the influence of its nearest 10 neighbors by 0.9) and renormalized from 0 to 1; this procedure was repeated 1500 times until the values at each iteration remained stable. The resulting vector was rank-ordered, and these ranks were considered the ranks for each glomerulus along a single axis that spanned across the medial glomerular domain.

Sister glomeruli are matching pairs between the medial and lateral glomerular domains. Linear regression was performed to evaluate how well the ranks of each medial glomerulus could be predicted based on the positions of its corresponding lateral sister, to assess the conservation of the glomerular maps in each domain. To do so, support vector machine regression models (with a rbf kernel and C=1 regularization) were trained using 5-fold cross-validation on the z-scored lateral positions to predict the glomerular ranks.

Symmetric glomeruli are those that were identified in both the left and right OB. For these glomeruli, we could also evaluate to what extent nearest-neighbor relationships were preserved across the two OB. For each OB, pairwise distances were calculated for all pairs of glomeruli within the same domain, and these distances were compared across OB.

### Comparisons of glomerular positions with OR genomic features

ORs were categorized as class I and class II based on their amino acid sequences. While class I ORs are located within a single genomic cluster on chromosome 7, class II ORs are located in genomic clusters that are scattered throughout the genome. ORs that were located on the same chromosome within 3 Mb of each other were defined as being within the same genomic cluster. Phylogenetic distances were calculated for pairs of OR, as previously described^34^. In brief, phylogenetic trees were constructed from OR amino acid sequences, and the cophenetic distance of the resulting pairwise phylogenetic distance matrix was taken to be the phylogenetic distance between pairs of ORs. The distribution of phylogenetic distances was evaluated for pairs of class II ORs whose glomeruli were located at given spatial distances within the same OB domain. Additionally, for each OR genomic cluster, the mean phylogenetic and glomerular distance were evaluated for the set of OR pairs whose ORs were both from that genomic cluster. As an example, the OR cluster at Chr10:128.9–130.2 Mb contains ORs that are both phylogenetically and spatially similar to each other, and this cluster is enriched for the set of ORs that express *Cd36*.

### Prediction of glomerular positions from OR transcriptomic features

Apart from genomic features, each OSN subtype is associated with a unique transcriptome, which can be summarized by the expression of a set of ∼1,300 highly-variable genes (HVGs) whose expression varies across mature OSNs. These HVGs, and their mean expression for OSN subtypes expressing given ORs were evaluated from a recent single-cell RNA-sequencing (scRNA-seq) dataset, as previously described^34^. Linear regression was performed for geometrically-confirmed glomeruli to evaluate to what extent HVG expression for each OSN subtype could predict its respective glomerular positions in each OB domain. A scikit-learn regression pipeline was constructed in which, on training data, HVG expression for each OSN subtype was z-scored, the top 500 HVGs were selected, and PCA was performed to reduce the dimensionality of this data to 200 dimensions. Elastic net linear regression models (l1_ratio=0.05, alpha=1) were fit on these PCs, and the performance of this pipeline was then evaluated on held-out test ORs. Predicted positions for each glomeruli were evaluated using leave-one-out cross-validation separately for each domain (medial and lateral) for each axis (anterior-posterior and dorsal-ventral). Additionally, the median absolute error was evaluated for each axis in each domain across 100 restarts in which 20% of glomeruli were randomly held out.

OSNs expressing different ORs are thought to express different levels of axon guidance genes, which could influence the targeting of their axons to their respective glomerular positions. To more directly test this hypothesis, linear regression was also performed using only the expression of each axon guidance gene, as measured via scRNA-seq, as features. Axon guidance genes were defined as genes representing known axon guidance or cell adhesion molecules, and which included 183 genes that contained the following prefixes: *Slit, Robo, Epha, Ephb, Efn, Nrp, Sema, Plx, Kirrel, Pcdh, Cdh, Tenm*. Linear regression was performed as described above using a similar pipeline, except the gene feature selection step was omitted, only the top 50 PCs were kept, and the elastic net models were trained with an l1_ratio=0.01.

To group axon guidance and HVGs into sets of related genes, the predicted positions from the linear regression models were correlated with the observed positions for each domain and axis and the resulting correlation matrix was hierarchically clustered via scikit-learn’s AgglomerativeClustering (with k=7 clusters). To summarize the average expression of genes associated with each cluster, the gene expression for each gene was z-scored across glomeruli and then averaged for each cluster. For example, the resulting cluster 2 was enriched for genes whose expression was the highest in OSN subtypes whose glomeruli projected to the anterior OB, whereas the expression of genes in cluster 6 was the highest for OSN subtypes enriched in the medial region of the OB. For visualization purposes, the average gene expression for each OSN subtype was plotted on the OB maps at the locations of its associated glomeruli. A similar approach was also used to identify HVGs whose expressions across OSN subtypes was highly-correlated with glomerular positions.

### Gene expression analysis of deep layer structure

#### Rotating axis differential expression analysis

Sections 38 to 147 were subset from the full dataset for deep layer analysis to avoid potential artifacts from non-main OB regions like the accessory olfactory bulb (AOB) and anterior olfactory nucleus (AON). Capture spots from Leiden Cluster 1, Leiden Clusters 8 and 9, and Leiden Cluster 6 were isolated from the full dataset and analyzed based on their morphological alignment to the deep granule cell layer, superficial granule cell layer, and mitral cell layer (respectively). For each cluster/layer, the 110-section dataset was collapsed along the z-axis into a 2D plane.

The symmetrized right OB was reflected onto the left OB and axes were imposed onto the data at 10 prespecified rotation angles from the glomerular axis of symmetry (set at 0°) (i.e., 0°, 18°, 36°… 162°, 180°). For each angle, differential expression analysis (DEA) was performed using the Wilcoxon Rank Sum Test for capture spots distributed on each side of the axis, comparing the right (medial) portion of the OB to the left (lateral) portion. This analysis produced DEA with log_2_FC and adjusted p-value for each gene at each of the prespecified axis angles. A gene with a positive log_2_FC represents upregulation on the right side of the axis, and a gene with a negative log_2_FC represents upregulation on the left side of the axis. This output was then filtered to only axes with adjusted p-value less than 0.05. Then, for each gene, the analysis was filtered to retain the top axis angle that generated the DEA result with the largest absolute log_2_FC value. Finally, the gene/axis pairs with the top 50 largest absolute log_2_FC values were retained for visualization (**Table S2**).

## SUPPLEMENTS

**Figure S1:**
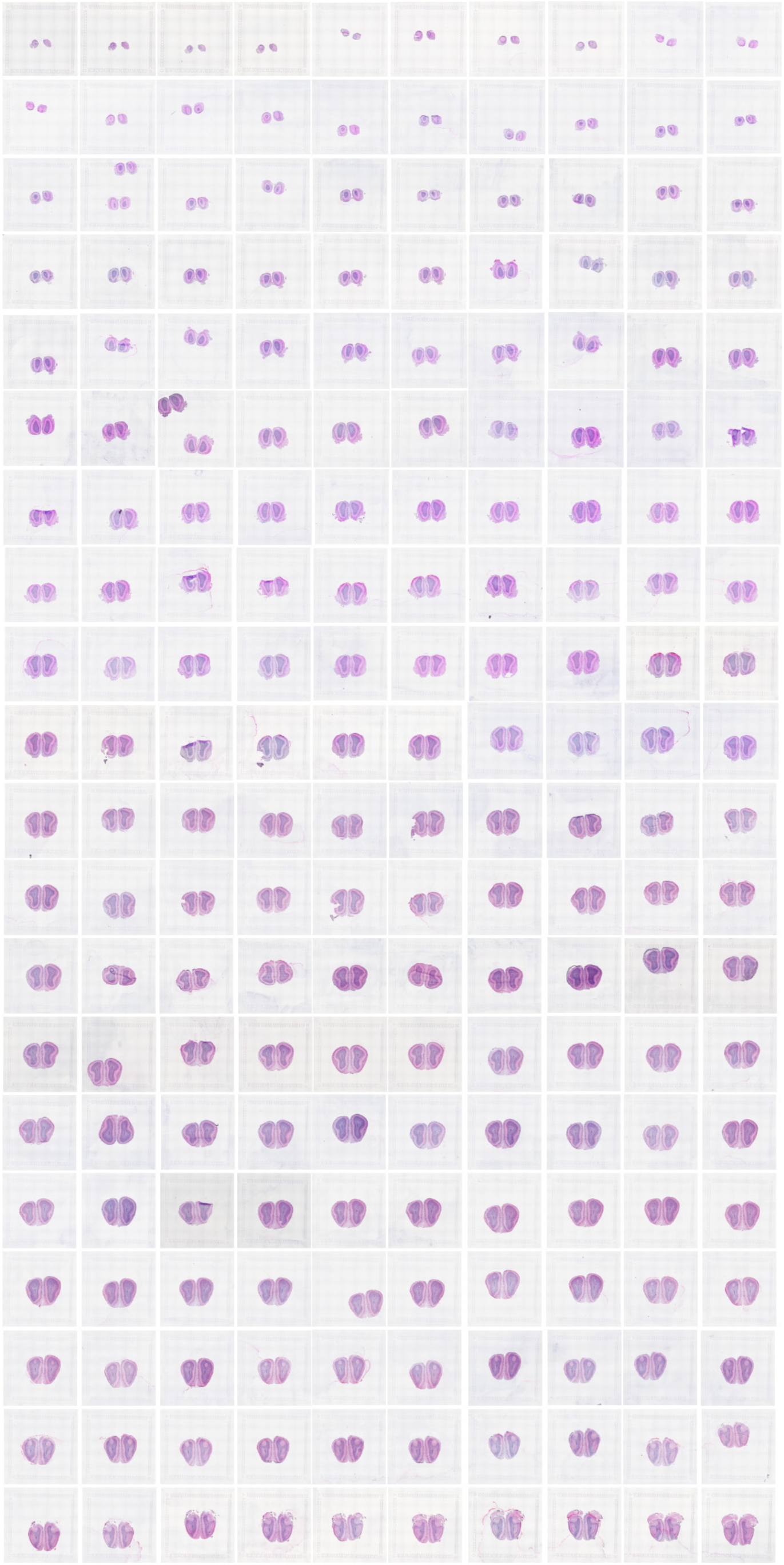
Morphology of the 200 consecutive sections of OB included in spatial transcriptomics dataset. Brightfield images of OB tissue sections mounted onto spatial transcriptomics microarrays. Sections were stained with hematoxylin and eosin for alignment of morphology and gene expression data.

**Figure S2:**
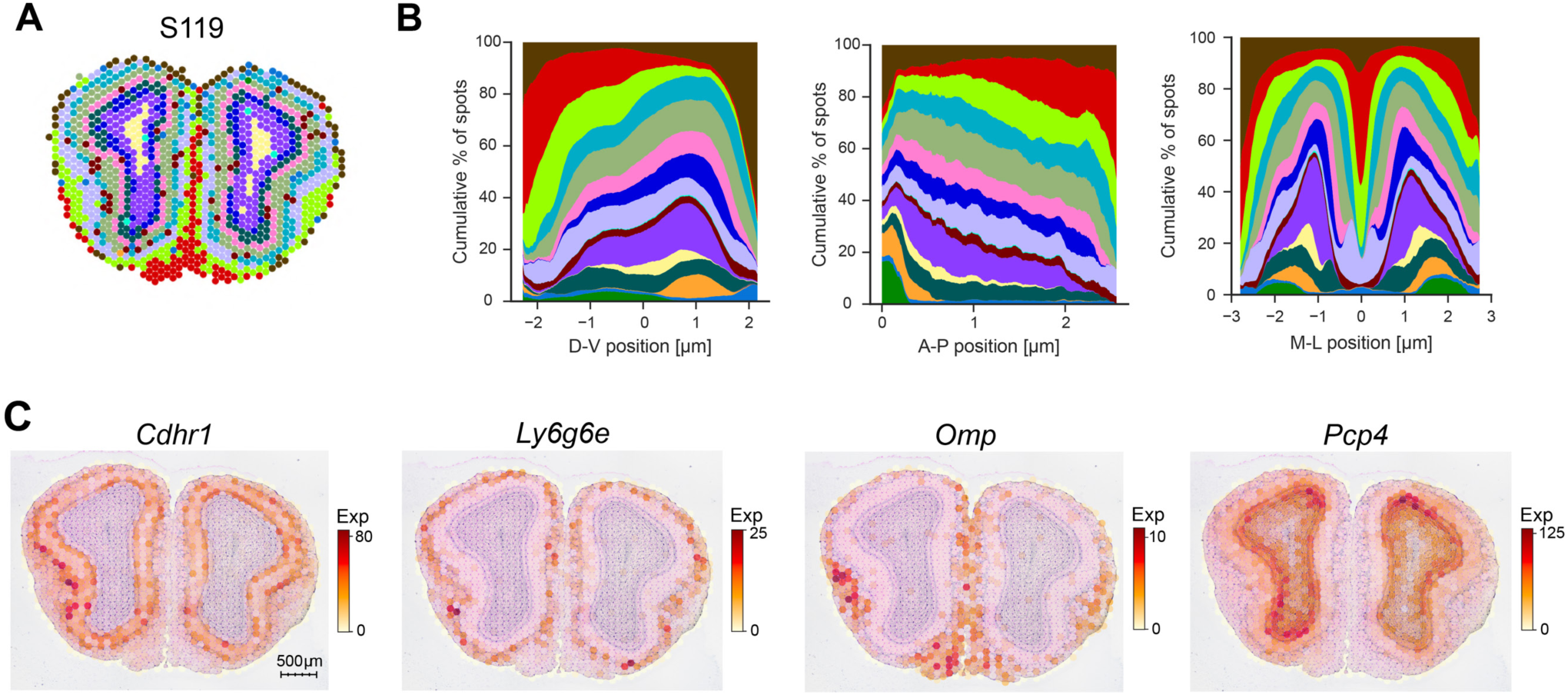
Leiden clustering of integrated capture spots across 200 sections. **(A)** Capture spots colored by Leiden cluster in section 119 out of 200 from the dataset. **(B)** Cumulative distribution of capture spots across Leiden clusters across the entire dataset along the D-V, A-P, and M-L axis. Color indicates cluster identity. Values were binned and smoothed across the X, Y, and Z axes. **(C)** Raw expression of known genetic markers for mitral and tufted cells (*Cdhr1*), external tufted cells (*Ly6g6e*), olfactory sensory neurons (*Omp*), and granule cells (*Pcp4*).

**Figure S3:**
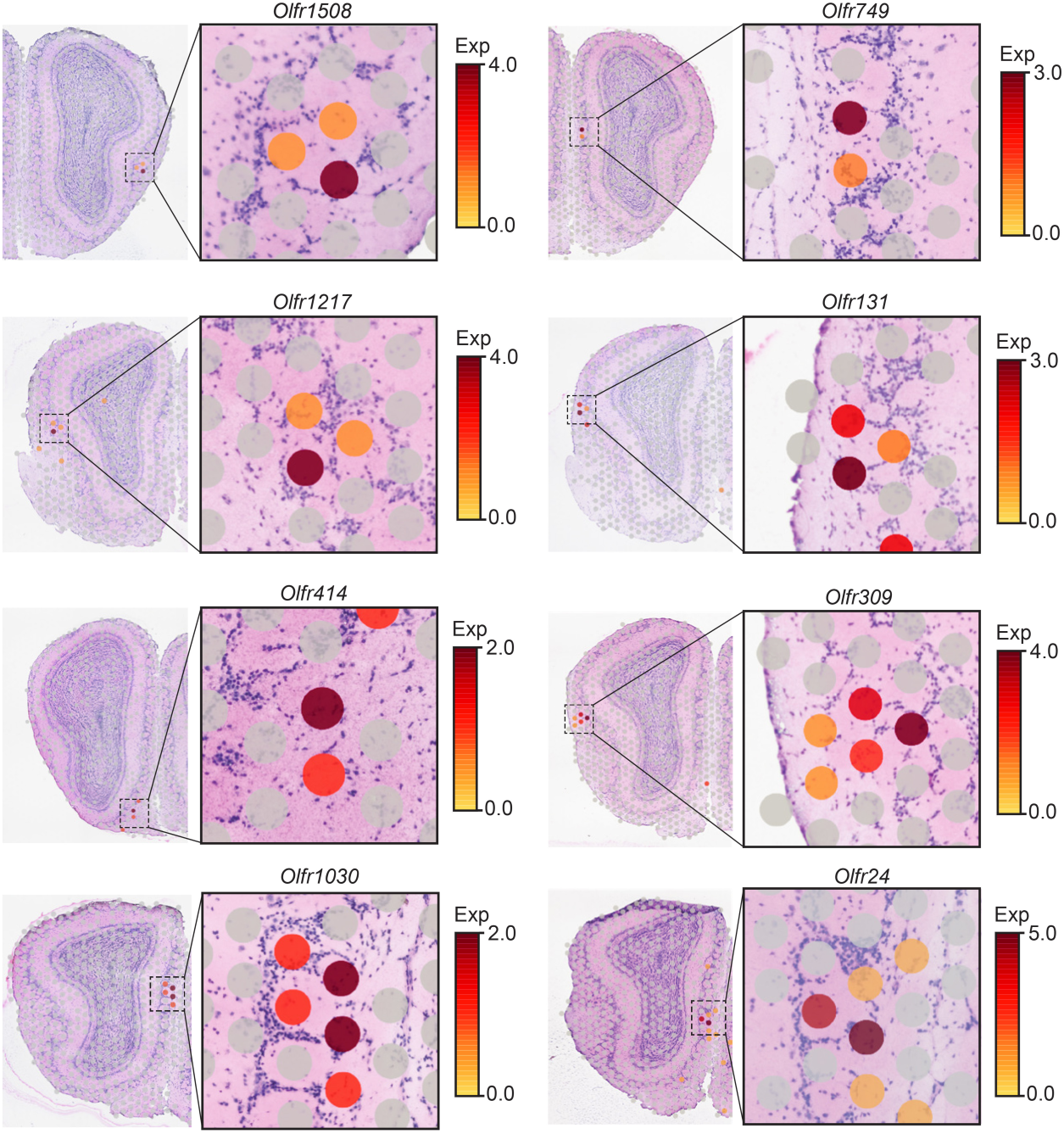
Odorant receptor expression in capture spots surrounding glomeruli. Examples of expression of *Olfr1508*, *Olfr749*, *Olfr1217*, *Olfr131*, *Olfr414*, *Olfr309*, *Olfr1030*, and *Olfr24* in individual tissue sections. High OR expression is restricted to 2-6 capture spots surrounding individual glomeruli.

**Figure S4:**
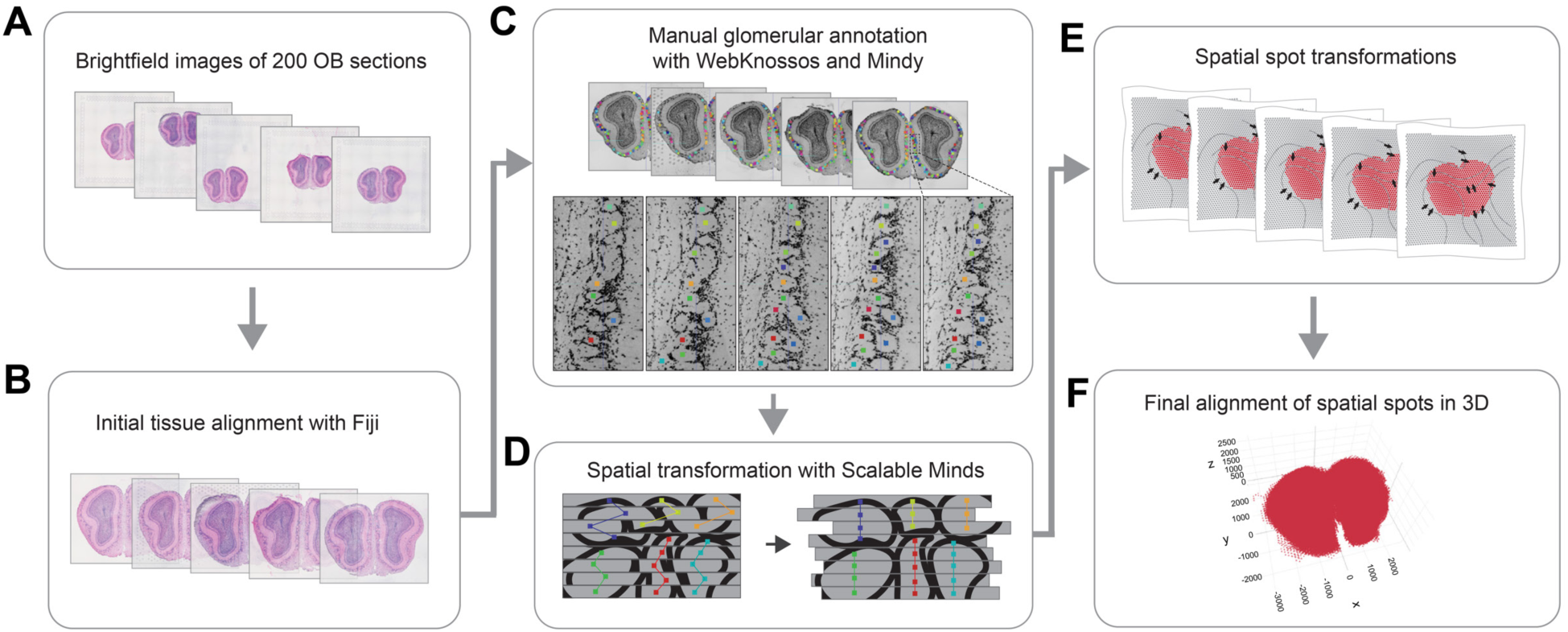
Morphological alignment of tissue sections to generate 3D map of OB. **(A)** The process of aligning spatial transcriptomics data from 200 consecutive OB sections involved the acquisition of brightfield images of OB sections stained with hematoxylin and eosin and initial tissue alignment of the sections with Fiji **(B)** (see Methods). **(C)** Further refinement of the morphological alignment was then performed by annotating glomeruli, where individual glomeruli were identified across consecutive sections, using the nuclei of the periglomerular layer to distinguish glomeruli. Colored squares denote the centers of glomeruli that were tracked. Glomeruli were annotated using WebKnossos, and the annotation of over 2,500 glomeruli was performed using the annotation service Mindy. **(D)** Spatial transformations were then performed with Scalable Minds to align sections based on the annotations. **(E)** Finally, the spatial coordinates of the capture spots from the spatial transcriptomics data were transformed according to the morphological transformations, resulting in a final 3D plot of aligned capture spots from all 200 sections **(F)**.

**Figure S5:**
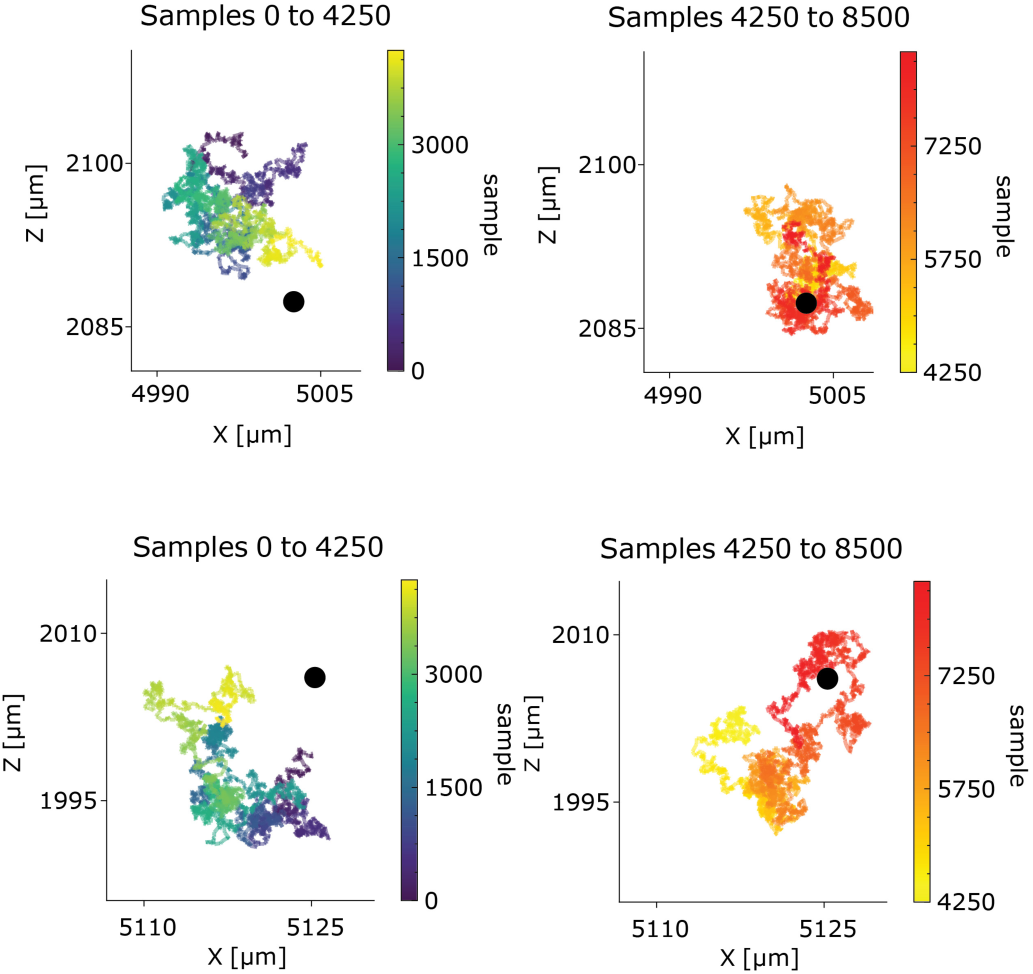
Predicted glomerular position across successive iterations of the MCMC algorithm for two different glomeruli. The predicted position of a glomerulus for iterations (‘Samples’) 0 to 4250 (Left) and iterations 4250 to 8500 (Right), with color in the plot indicating iteration. The black dot in both plots is the final predicted glomerular position.

**Figure S6:**
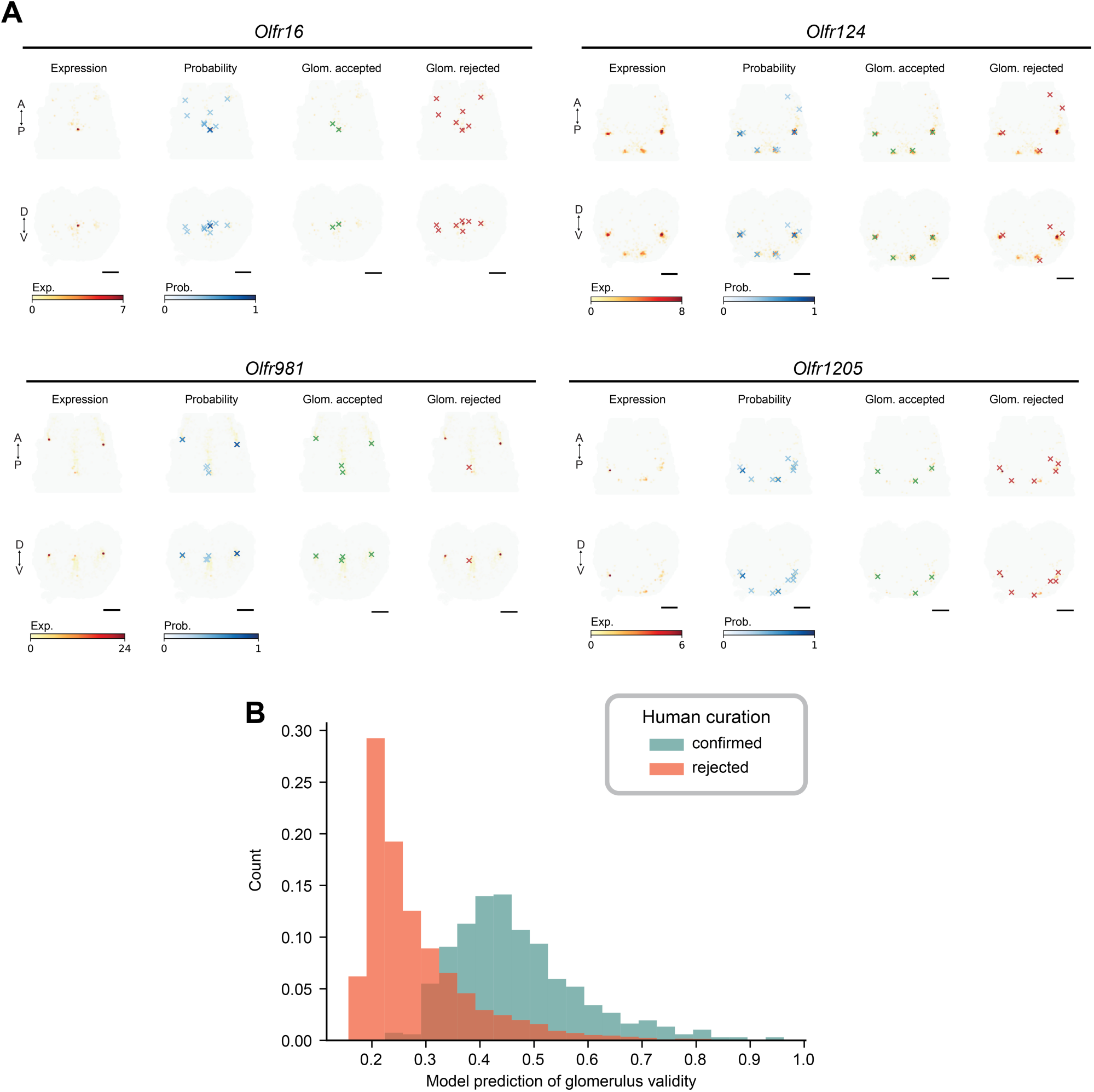
Human expert curation of glomerular positions output by PGP model. **(A)** Examples of accepted and rejected glomeruli for *Olfr16*, *Olfr124*, *Olfr981*, and *Olfr1205* after human curation. First column shows raw expression, and second column shows glomerular positions output by the PGP model (Xs) colored by probability from 0 to 1. Third and fourth columns show positions accepted (green) or rejected (red) during expert curation. **(B)** Comparison of model-generated probability values for glomerulus validity and human curation decisions. Confirmed glomeruli (green) and rejected glomeruli (red) showed good alignment with the model’s goodness of fit for putative glomerular locations.

**Figure S7:**
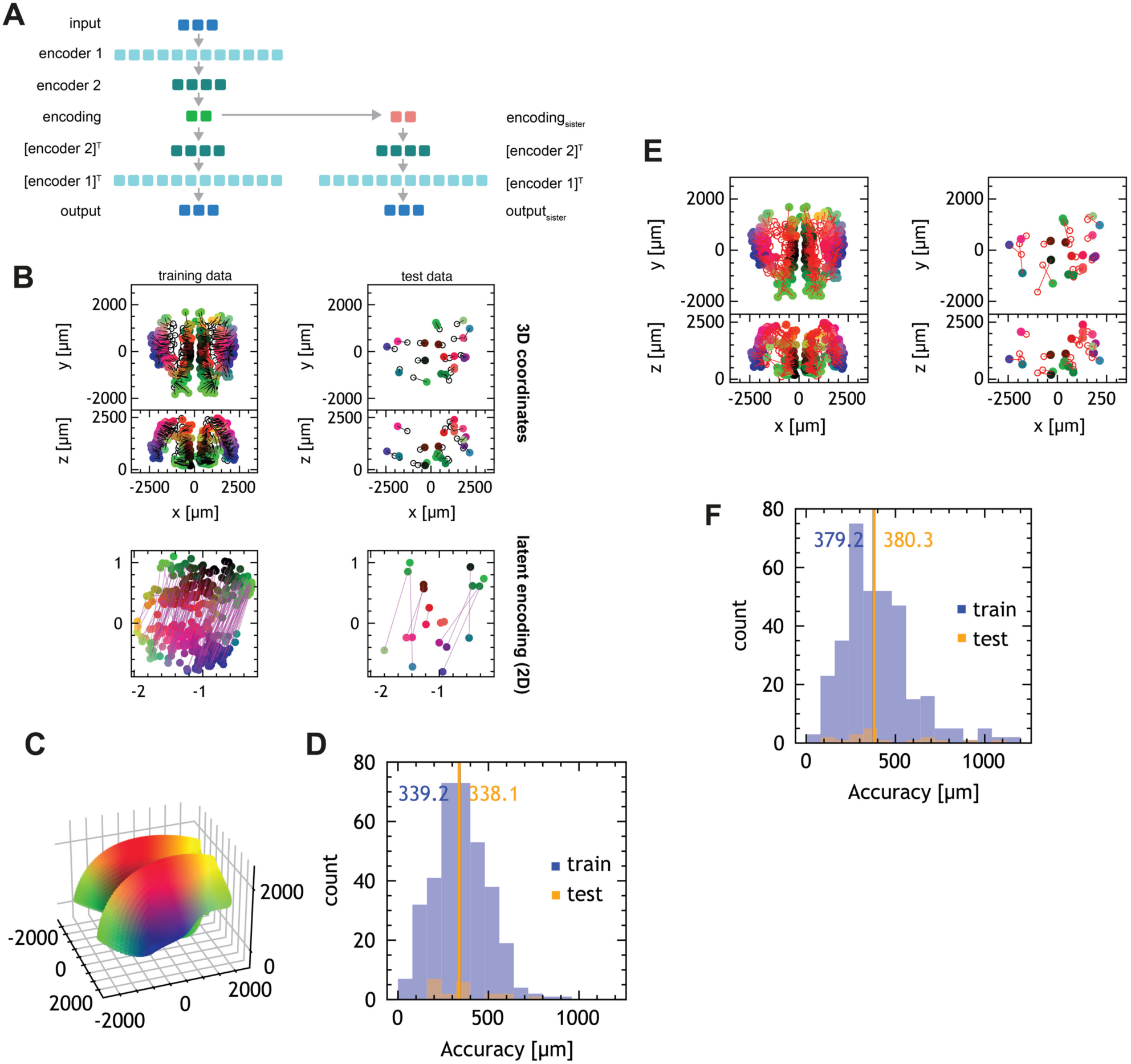
Dual autoencoder model for mapping glomeruli and predicting sister glomerulus locations. **(A)** Architecture of the dual autoencoder model. After symmetrization, the glomerular coordinates are encoded through two encoder layers into a two-dimensional latent representation. The latent representation is decoded via the transposition of the same autoencoder layers to reconstruct the original data. A second branch of the model predicts the encoded location of sister glomeruli with a linear layer from the encoded glomerular location. Sister glomeruli are decoded with the same layers into 3D coordinates. **(B)** Results after training of the dual autoencoder. Filled circles represent the original coordinates, and the linked open circles represent the corresponding reconstructed position. The color code is identical between the original coordinate plot (top) and the 2D latent map (bottom). In the latent map, sister glomeruli are shown as magenta lines. **(C)** The 3D manifold learned by the autoencoder, showing 3D locations corresponding to each part of the latent space. **(D)** Reconstruction precision after encoding and decoding. The lines represent mean precision for train and test data. **(E)** 3D locations of test and train data for sister glomerulus prediction. **(F)** Predictive accuracy of sister glomerulus precision of the dual autoencoder. Sister glomerulus locations can be predicted up to 380.3 µm on previously unseen data, defining an upper bound for the precision of the glomerular map.

**Figure S8:**
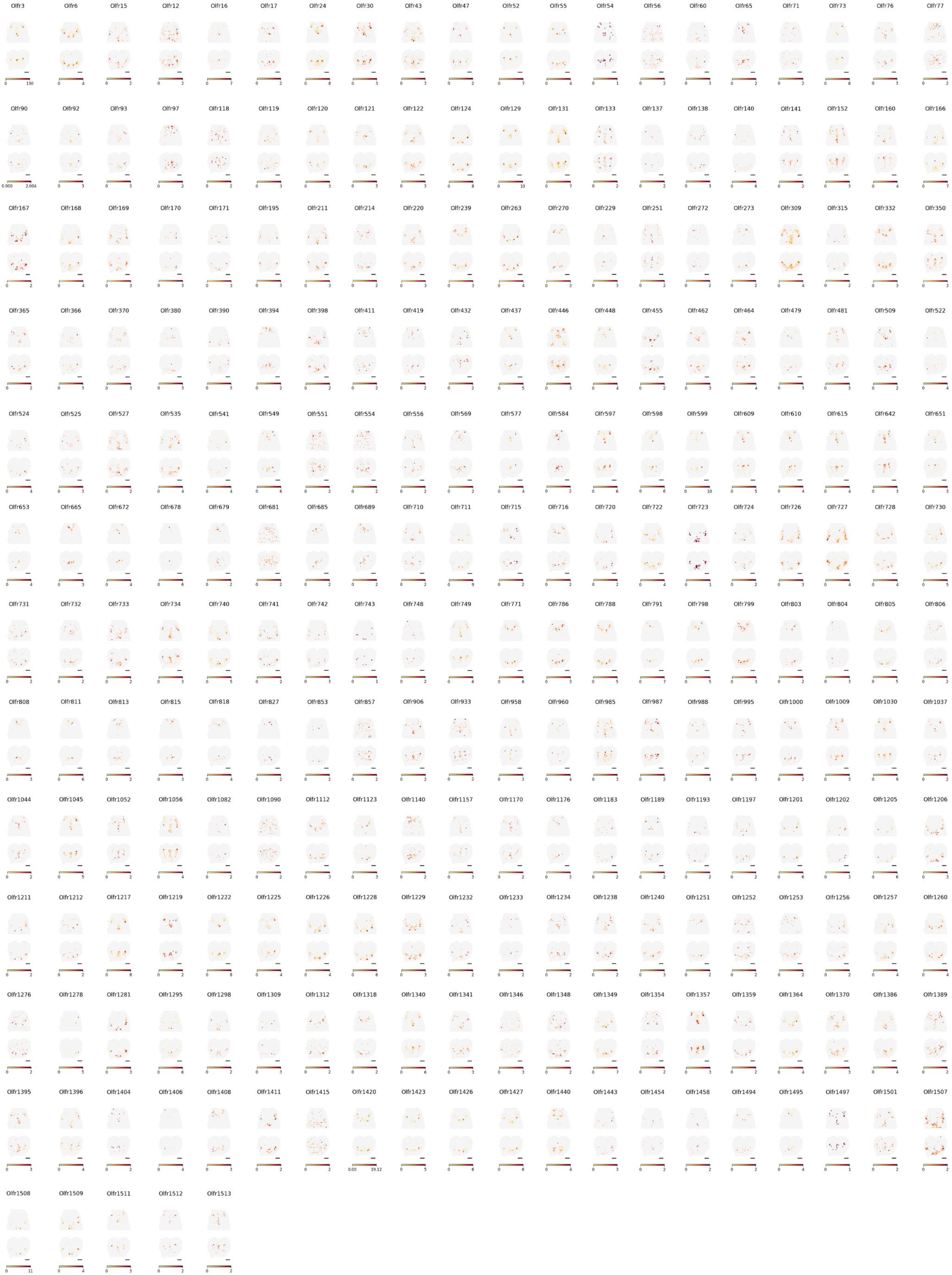
Expression of OR genes included in the final dataset of 968 glomeruli. Raw expression of 255 OR genes, visualized as a maximum Y-projection (top) and Z-projection (bottom).

**Figure S9:**
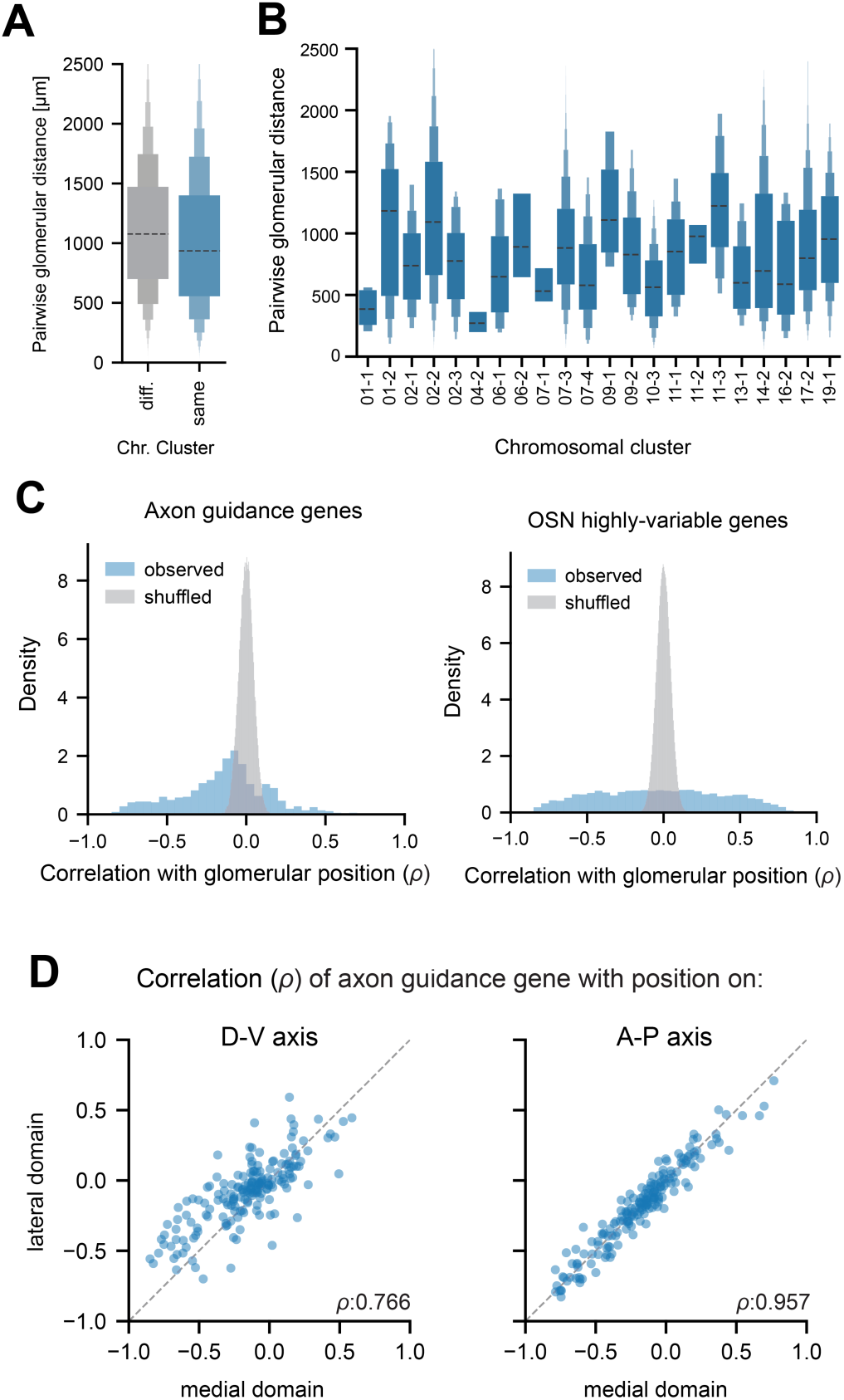
OR identity and transcriptome-wide gene expression correlate with glomerular position. **(A)** Boxplot of pairwise glomerular distance for glomeruli expressing ORs on the same or different chromosomal cluster. **(B)** Boxplot of pairwise glomerular distance for ORs in 22 chromosomal clusters. **(C)** Histograms of observed correlation with glomerular position for axon guidance genes (left) and highly variable genes (right plot) versus shuffled control. **(D)** Correlations of predicted positions of glomeruli using a regression model trained on the axon guidance gene set. The model performed better at predicting positions of glomeruli across the A-P axis than the D-V axis (*ρ*=0.957 and 0.766, respectively).

**Figure S10:**
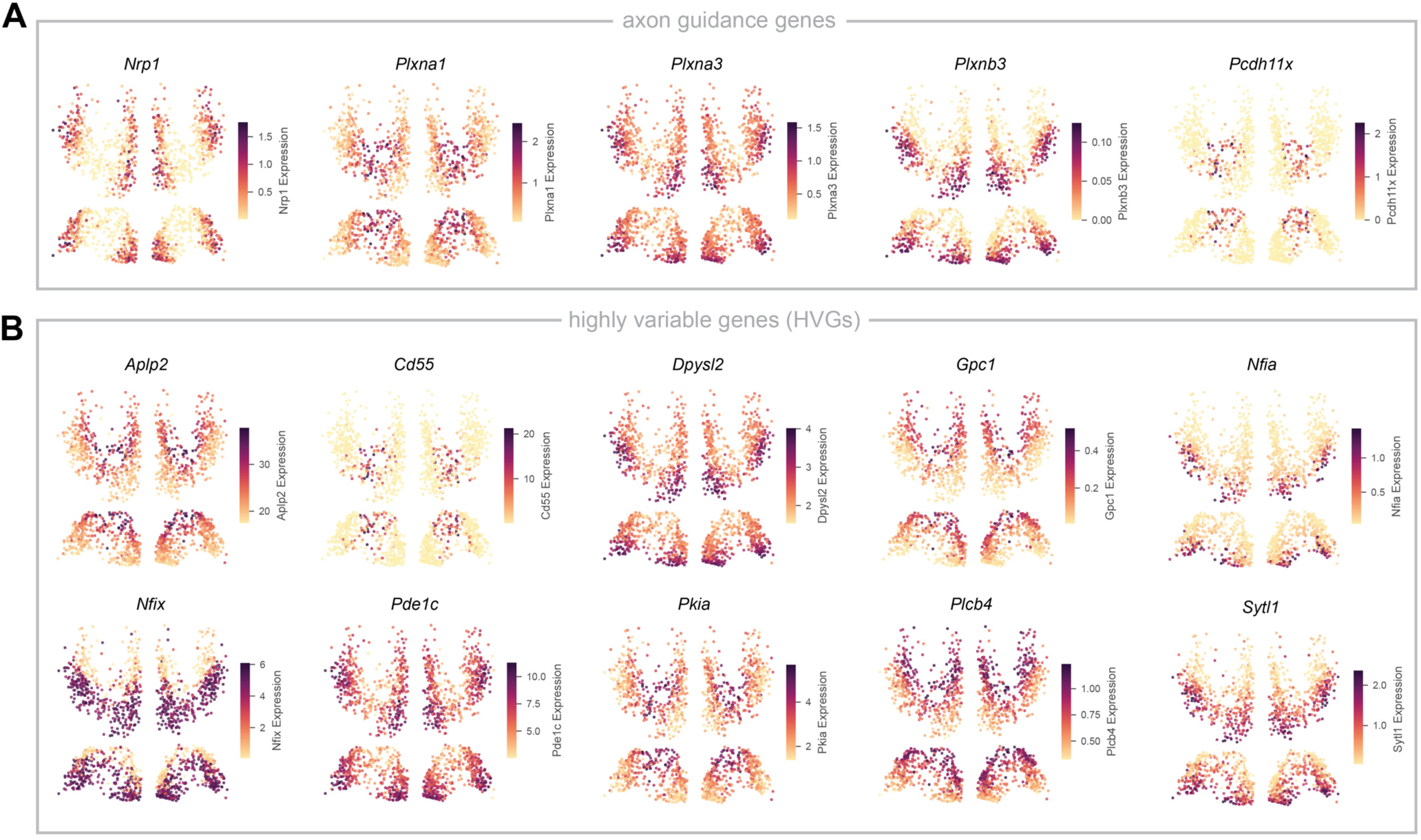
Additional expression examples of axon guidance genes and HVGs. **(A)** Expression of individual axon guidance genes *Nrp1*, *Plxna1*, *Plxna3*, *Plxnb3*, and *Pcdh11x* the in glomerular map. The top plot shows Z-projection and bottom plot shows Y-projection of the OB. **(B)** Expression of highly variable genes *Aplp2*, *Cd55*, *Dpsyl2*, *Gpc1*, *Nfia*, *Nfix*, *Pde1c*, *Pkia*, *Plcb4*, and *Sylt1* in the glomerular map. Top plot shows Z-projection and bottom plot shows Y-projection of the OB.

**Fig S11:**
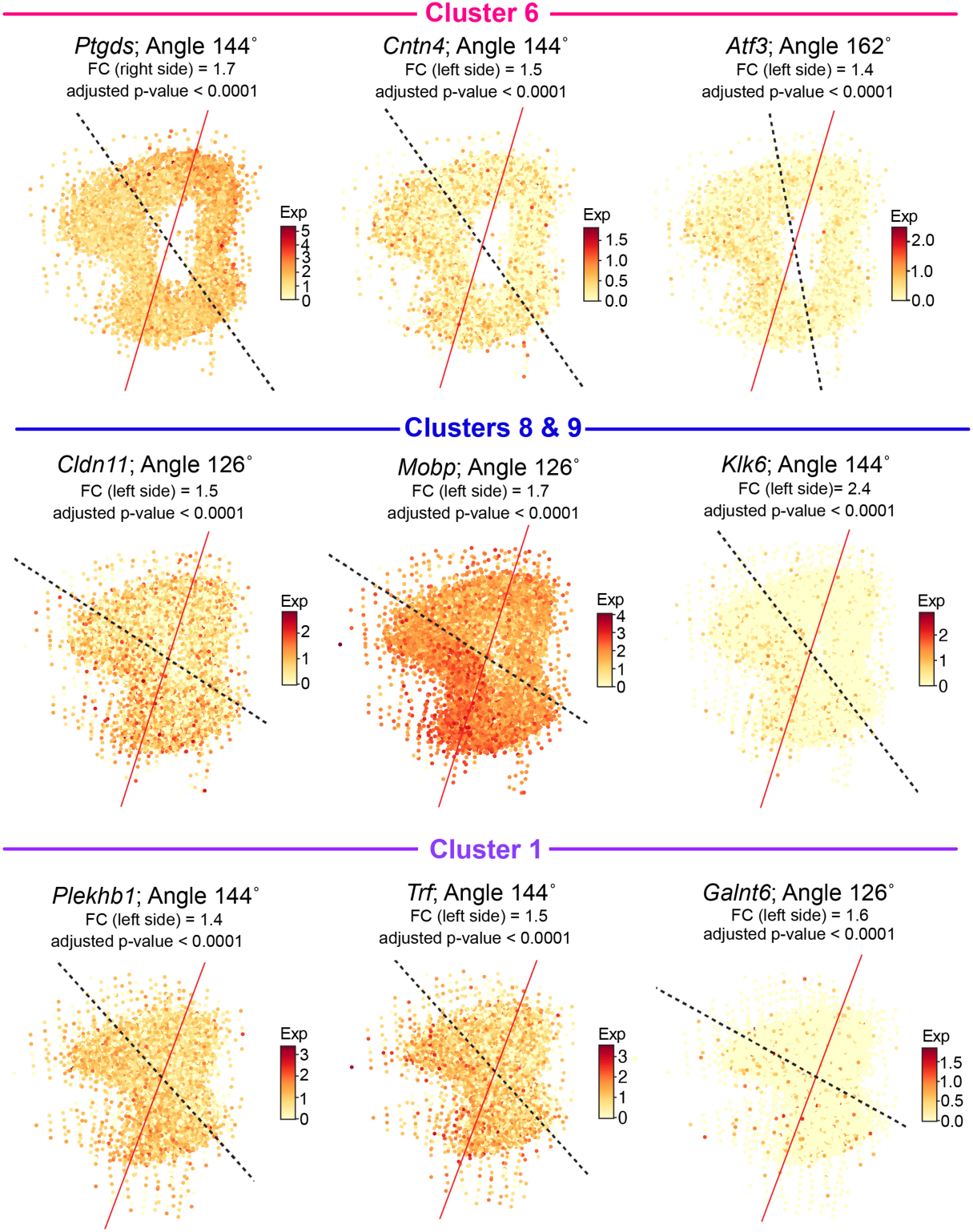
Individual gene expression across alternative axes to the glomerular axis of symmetry. Expression of 9 genes (*Ptgds*, *Cntn4*, *Atf3*, *Cldn11*, *Mobp*, *Klk6*, *Pelkhb1*, *Trf*, *Galnt6*) exhibiting differential expression across a given axis, in black (in degrees from the glomerular axis, in red). Expression shown is from 110 sections flattened in 2D and isolated by Leiden cluster (see also Methods).

